# Differential prolyl hydroxylation by six Physcomitrella prolyl-4 hydroxylases

**DOI:** 10.1101/2024.03.26.586753

**Authors:** Christine Rempfer, Sebastian N. W. Hoernstein, Nico van Gessel, Andreas W. Graf, Roxane P. Spiegelhalder, Anne Bertolini, Lennard L. Bohlender, Juliana Parsons, Eva L. Decker, Ralf Reski

## Abstract

The hydroxylation of proline residues to 4-trans-hydroxyproline (Hyp) is a common post-translational protein modification in plants, mediated by prolyl 4-hydroxylases (P4Hs). Hyps predominantly occur in a group of cell wall proteins, the Hydroxyproline-rich glycoproteins (HRGPs), where they are frequently *O*-glycosylated. While prolyl-hydroxylation and *O*-glycosylation are important, e.g. for cell wall stability, they are undesired on plant-made pharmaceuticals. Sequence motifs recognized for prolyl-hydroxylation derived from vascular plants were proposed but did not include data from mosses, such as Physcomitrella. Here, a phylogenetic reconstruction of plant P4Hs identified six P4Hs in four subfamilies in mosses. We analysed the amino acid sequences and structural environments around Hyps in Physcomitrella utilizing 73 Hyp sites in 24 secretory proteins from multiple MS/MS datasets, and found that prolines in close proximity to other prolines, alanine, serine, threonine and valine were preferentially hydroxylated. About 95% of the Hyp sites were predictable with a combination of previously defined motifs and methods. In our data, AOV (Ala-Hyp-Val) was the most frequent prolyl-hydroxylation pattern. Additionally, short arabinose chains were attached to Hyps in two cell-wall pectinesterases. A combination of 443 AlphaFold structure models and our MS data of peptides with nearly 3000 proline sites found Hyps predominantly on protein surfaces in disordered regions. Moss-produced human erythropoietin (EPO) exhibited plant-specific *O*-glycosylation with arabinose chains on two Hyps. This modification was significantly reduced in a P4H1 single knock-out (KO) Physcomitrella mutant. Quantitative proteomics after isotope labelling with different P4H-KO moss mutants revealed specific changes in the amount of proteins, including HRGPs, and modified prolyl-hydroxylation pattern from the mutants, suggesting a differential function of the six Physcomitrella P4Hs. Quantitative RT-PCR proved a differential effect of single P4H KOs on the expression of the other five *p4h* genes, suggesting a partial compensation of the mutation. AlphaFold-Multimer models for Physcomitrella P4H1 and its target EPO peptide superposed with the crystal structure of Chlamydomonas P4H1 and a peptide substrate suggested significant amino acids in the active centre of the enzyme that form H-bonds with the peptide substrate, and revealed differences between P4H1 and the other five Physcomitrella P4Hs.

**Highlights:** Mosses, such as Physcomitrella, encode six prolyl 4-hydroxylases in four subfamilies Computational analysis reveals specific environment for proline hydroxylation AlphaFold structure models and mass spectrometry reveal features of modification Quantitative proteomics and mutant analysis identify relevant modifying enzyme Modelling suggests relevant amino acids in active centre of the modifying enzyme

## 1. Introduction

Hydroxyproline-rich glycoproteins (HRGPs) are important structural proteins of the plant cell wall with functions in growth, stress response, signalling and reproductive development (Cassab and Varner, 1988; Han et al., 2001; Deepak et al., 2010; Draeger et al., 2015). HRGPs possess a signal peptide that mediates their entry into the endoplasmic reticulum (ER). In the ER and Golgi apparatus, proline residues of HRGPs are frequently hydroxylated by prolyl-4 hydroxylases (P4Hs) to 4-trans hydroxyproline (Hyp) that serves as anchor for the attachment of *O*-glycans. In a final step the proteins are secreted to the apoplast (Silva et al., 2020; Mishler-Elmore et al., 2021) where HRGPs cross-link with other elements of the cell wall *via* their glycans (Fruleux et al., 2019).

One class of HRGPs are extensins that contain characteristic hydrophilic Ser-(Pro)_n≥2_ motifs in which the prolines are hydroxylated and *O-*glycosylated with linear chains of one to five arabinoses (Hijazi et al., 2014; Kieliszewski and Lamport, 1994). *O*-glycosylated Hyps are associated with an increased stability of polyproline-II helical conformations as they exist in extensins (Owens et al., 2010; Marzol et al., 2018; van Holst and Varner 1984). Extensins undergo cross-linking mediated by hydrophobic tyrosine-rich motifs that *in vitro* depend on the presence of arabinosylated Hyps (Chen et al., 2015).

Another class of HRGPs are arabinogalactan proteins (AGPs) that carry large, branched, and complex glycans with variable structures which constitute up to 90 % of the glycoprotein mass. Glycans on AGPs consist mainly of galactose and arabinose, but also rhamnose, fucose and glucuronic acid (Silva et al., 2020). Characteristic for AGPs is a high proportion of the amino acids (AAs) proline, alanine, serine and threonine (PAST), that facilitate the bioinformatic identification of AGPs using thresholds with PAST contents of 50% and above (Schultz et al., 2002; Ma and Zhao, 2010; Showalter et al., 2010; Johnson et al., 2017). Target sites of *O*-glycosylation on AGPs are repeats of Ala-Hyp, Ser-Hyp, Thr-Hyp and Val-Hyp that, when present in synthetic peptides, showed varying degrees of prolyl-hydroxylation and *O*-glycosylation depending on the neighbouring AAs (Shpak et al., 1999; Tan et al., 2003).

Based on different studies on prolyl-hydroxylation motifs in HRGPs (e.g., Shpak et al., 1999; Shpak et al., 2001), Gomord et al. (2010) proposed a motif for prolyl-hydroxylation and subsequent *O*-glycosylation, the [Ala ⁄ Ser ⁄ Thr ⁄ Val]-Pro_(1,4)_-X_(0,10)_-[Ala ⁄ Ser ⁄ Thr ⁄ Val]-Pro_(1,4)_ glycomodule, where X can be any AA. According to the Hyp contiguity hypothesis mostly arabinose chains are attached to blocks with neighbouring Hyps, probably due to space constraints, while arabinogalactans are added more often to single non-contiguous Hyp residues (Kieliszewski and Lamport, 1994). Subsequently, Canut et al. (2016) proposed an extended prolyl-hydroxylation code including data from 25 plant species taking into consideration the neighbouring AAs of Hyps. However, multiple exceptions from these rules are known. For example, in a *Lolium multiflorum* AGP only the first proline in the motif Ser-Pro-Pro-Ala was hydroxylated even though both prolines should be hydroxylated according to the extended prolyl-hydroxylation code (Canut et al., 2016). More recently, an R package for analysis of HRGPs was developed including a function for the prediction of Hyp sites in plant proteins (Dragićević et al., 2020). It is based on a machine-learning algorithm that was trained on plant protein sequences from UniProt, some of them containing experimentally determined Hyp sites.

Usually, plants have a set of P4H isoforms with diverging substrate preferences. For example, *Arabidopsis thaliana* (Arabidopsis) P4H1 hydroxylates mostly the second proline in the Pro-Pro-Gly motif from collagen, which is a substrate for mammalian P4Hs (Gorres and Raines, 2010), whereas hydroxylation of such peptides is inefficient for P4H2 (Hieta and Myllyharju, 2002; Tiainen et al., 2005). Comparably, in *Nicotiana benthamiana* (Nicotiana) three of four examined P4Hs from different branches of the phylogenetic tree were able to hydroxylate a proline in an EPO peptide (Mócsai et al., 2021). The same proline is also prolyl-hydroxylated in recombinant EPO from Physcomitrella (Weise et al., 2007), specifically by P4H1 (Parsons et al., 2013).

Prolyl-hydroxylation and subsequent *O*-glycosylation on recombinant proteins can deteriorate plant-made pharmaceuticals (PMPs) because mammals may show an immune response against the arabinose residues of plant-specific *O*-glycans (Altmann, 2007). Knowledge of preferred prolyl-hydroxylation motifs of the different P4H isoforms in plants would be of great help to solve the issue of undesired prolyl-hydroxylation and Hyp-anchored *O-*glycosylation on PMPs. The production of PMPs relies on vascular plants, such as Nicotiana, and on the moss Physcomitrella. While the general *N*-glycosylation pattern is similar between the two (Koprivova et al., 2003; Decker et al., 2014), there are subtle differences (Stenitzer et al., 2022), suggesting that the existing prediction tools for prolyl-hydroxylation and *O*-glycosylation might not be sufficient for the prediction in Physcomitrella.

Here, we studied preferred prolyl-hydroxylation patterns in the model moss Physcomitrella (Lueth and Reski, 2023) that is an attractive production system for PMPs (Reski et al., 2015, Decker and Reski, 2020) and combines advantages such as high gene-targeting rates via homologous recombination, facilitating precise genome engineering (Hohe et al., 2004; Reski et al., 2018; Chen et al., 2024), a fully sequenced genome, and a published secretome (Lang et al., 2016; Lang et al., 2018; Hoernstein et al., 2018).

## 2. Methods

### 2.1 Phylogenetic reconstruction

With the V3.3 Physcomitrella P4H protein sequences as queries, BLASTp-like searches were performed with DIAMOND (version 2.1.8; Buchfink et al., 2021) in ‘ultra-sensitive’ mode against the protein sequences of 18 selected plant species: *Anthoceros angustus* (Zhang et al., 2020), *Arabidopsis thaliana* (Cheng et al., 2017), *Calohypnum plumiforme* (Mao et al., 2020), *Ceratodon purpureus* (Carey et al., 2021), *Ceratopteris richardii* (Marchant et al., 2022), *Funaria hygrometrica* (Kirbis et al., 2020), *Klebsormidium nitens* (Hori et al., 2014), *Marchantia polymorpha* (Bowman et al., 2017), *Mesotaenium endlicherianum* (Cheng et al., 2019), *Nicotiana benthamiana* (Ranawaka et al., 2023), *Oryza sativa* (Ouyang et al., 2007), *Populus trichocarpa* (Tuskan et al., 2006), *Selaginella moellendorffii* (Banks et al., 2011), *Sphagnum fallax* (Healey et al., 2023), *Sphagnum magellanicum* (Healey et al., 2023), *Spirogloea muscicola* (Cheng et al., 2019), *Takakia lepidozioides* (Hu et al., 2023) and *Vitis vinifera* (Jaillon et al., 2007). Initial hits were searched for signatures of the ‘2OG-Fe(II) oxygenase superfamily’ from Pfam (PF13640), the ‘PROLYL 4-HYDROXYLASE ALPHA SUBUNIT’ from PANTHER (PTHR10869) and the ‘Fe(2+) 2-oxoglutarate dioxygenase domain profile’ from prosite (PS51471) with InterProScan (version 5.66-98.0; Jones et al., 2014). Only sequences with all three signatures were kept and complemented by the P4Hs of *Homo sapiens* (UniProt P13674, O15460, Q7Z4N8, Q9NXG6) and *Mus musculus* (UniProt Q60715, Q60716, Q6W3F0, Q8BG58) as an outgroup for phylogenetic reconstruction. Several iterations of multiple sequence alignments and maxim-likelihood tree inferences were performed using MAFFT (version 7.149b; Katoh and Standley, 2013) and FastTree (version 2.1.11; Price et al., 2010), to filter out non-homologous sequences. The multiple sequence alignment of resulting protein sequences was performed with UPP/SEPP (version 4.5.2; Nguyen et al., 2015) and converted into a codon-aware alignment of associated coding sequences with PAL2NAL (version 14; Suyama et al., 2006). A maximum-likelihood tree was calculated with RAxML-NG (version 1.2.1; Kozlov et al., 2019) using the ‘GTR+G’ model and 1000 bootstrap replicates, outgroup-rooted and visualized using R (version 4.3.2; R Core Team, 2024) and the ggtree package (version 3.10.0; Yu et al., 2017).

### 2.2 Mass spectrometry data sets

Hyp sites were selected from a collection of MS/MS measurements using samples with an enrichment of secretory Physcomitrella proteins from ER, Golgi apparatus, cell wall and extracellular space. The datasets contained: preliminary measurements of an EPO-producing line (174.16), quantitative MS data from mixtures of EPO-producing maternal line (174.16; labelled with ^15^N as described in Mueller et al., 2014) with single KO lines of P4H1 – P4H6, respectively, separate measurements of EPO-producing maternal line (174.16) and P4H1 KO line, new Mascot searches of the published host cell proteome dataset (Hoernstein et al., 2018) using Hyp with zero to three arabinoses as variable modifications, Mascot search with no protease specificity from chemically deglycosylated protein samples digested with thermolysin, elastase or trypsin, and double KO of two galactosyltransferases (Δ*galt*2/3; Parsons et al., 2012).

### 2.3 Cultivation and isotopic labelling

Physcomitrella plants were grown in axenic suspension culture in 500 mL flasks containing 180 mL of Knop ME medium (Egener et al., 2002) in stable conditions of 22° C and a light cycle of 16/8 h at an intensity of 55 µmol/m^2^ s. The flasks were kept on a rotary shaker at 125 rpm and held on protonema stage by dispersion with an Ultra-Turrax (IKA, Staufen, Germany) at 18,000 rpm for 1 min every two weeks. After dispersion, the protonema was harvested with a 100 µm sieve and transferred into a 500 mL flask containing fresh Knop ME medium using sterile tweezers. After turraxation, sterility controls were taken directly from the Ultra-Turrax tip for each cell line to check for potential contamination according to the protocol from Heck et al. (2020). For the quantitative approach in the MS analysis, the maternal line was inoculated in a small amount of 5-10 gametophores in medium containing the heavy ^15^N isotope for at least 6 weeks with weekly turraxation.

For sample preparation, protonema suspension cultures were cultivated for one week in 400 mL BM medium (Parsons et al., 2012) in a 1 L aerated round flask (modified after Lang et al., 2011). The whole supernatant of each flask was precipitated with 10% TCA overnight on ice in a cold room (6 °C). Protein pellets were obtained by centrifugation at 15,000 x g and 2°C for 2 h. The pellet was resuspended in ice-cold acetone, transferred into a 50 mL tube and centrifuged at 5,000 x g 4°C for 15 min each time. The pellet was again resuspended in ice-cold acetone, transferred into a 1.5 mL tube and centrifuged at 14,000 x g, 4°C for 15 min. The supernatant was discarded and the pellet was air-dried.

The protonema was filtered using a 100 µm sieve and resuspended in extraction buffer (408 mM NaCl, 60 mM Na_2_HPO_4_ x 2 H_2_O, 10.56 mM KH_2_PO_4_, 60 mM EDTA, 1% plant protease Inhibitor cocktail (Sigma P9599), pH7.4, using 3 mL buffer / g protonema (fresh weight). The sample was then homogenized using an Ultra-Turrax for 10 min at 10,000 rpm on ice and subsequently centrifuged at 4500 x g for 3 min at 4°C. The supernatant was transferred into a new tube and again centrifuged at 4500 x g for 5 min at 4°C. The supernatant was precipitated with methanol/chloroform as described (Wessel and Flügge, 1984; Lang et al., 2011).

### 2.4 Chemical deglycosylation

Chemical deglycosylation was achieved with the GlycoProfile^TM^ Kit (Sigma Aldrich, PP0510) using 1 mg total protein, which was harvested from supernatant fractions as described above. All steps were performed as recommended by the manufacturer using the scavenging procedure. Subsequently, the sample was precipitated with acetone as described in Hoernstein et al. (2018).

### 2.5 In-solution digest of protein samples

In-solution digest of protein samples for MS analysis was modified after Reimann et al. (2020). In brief, each one pellet of precipitated total protein was dissolved in 100 µL 8 M urea and 50 mM ammonium bicarbonate (AmBic, Witney, UK). Protein concentration was determined either with a BCA assay (Pierce^TM^) or via absorbance measurement at 280 nm (A280). In the latter case, values were adjusted using an empirical correction factor according to Pace et al. (1995). A total amount of 20 µg total protein at a concentration of 1 µg/µL was employed for trypsin digestion. Heavy labelled (^15^N) and corresponding light labelled (^14^N) protein samples were mixed in a 1:1 ratio. Prior to digestion, samples were reduced and alkylated at a final concentration of 5 mM TCEP (37° C, 30 min) and 50 mM iodoacetamide (RT, 30 min in darkness). The reaction was quenched at a final concentration of 20 mM DTT and the sample solution was diluted with 50 mM AmBic to reach a final urea concentration of 2 M. Trypsin (V5117, Promega, Walldorf, Germany) was added at a ratio of 1:50 and the digestion was performed over night at 37° C. The chemically deglycosylated samples were digested with thermolysin at a ratio of 1:50 for 60 min at 45° C or with trypsin or elastase at 37°C respectively. Generated peptides were purified via C18 STAGE-Tips as described by Hoernstein et al. (2018) and eluted in 30 % ACN in 0.1% FA.

### 2.6 Mass spectrometry and data analysis

MS analysis of samples from the in-solution digests of the chemically deglycosylated samples was performed according to Top et al. (2019). Samples of the metabolically labelled samples were measured in the same way but using a 3 h gradient. Raw data were processed with Mascot Distiller 2.8.3.0 (Matrix Science, London, UK) and database searches were performed with Mascot Server (V2.7.0). Processed peak lists were searched against a database containing all Physcomitrella protein models V3 (Lang et al., 2018) and the sequence of human EPO (P01588). “15N Metabolic [MD]” was specified as quantitation option and carbamidomethylation (C + 57.021464 Da) was specified as fixed modification. Variable modifications were Gln → pyro Glu (N term Q − 17.026549 Da), oxidation (M + 15.994915 Da), hydroxyproline (P + 15.994915 Da), mono-arabinosylation (P + 148.037173 Da), di-arabinosylation (P + 280.079432 Da) and tri-arabinosylation (P + 412.121691 Da), deamidation (N + 0.984016 Da) and glycosylation (S + 162.052823 Da).

### 2.7 MS sample preparation

Identification of hydroxyproline sites or arabinosylated hydroxyproline residues was performed on human EPO recombinantly produced in Physcomitrella. In brief, cell culture supernatant of an EPO producing line (174.16; IMSC40216, www.moss-stock-center.org) (Weise et al., 2007; Parsons et al., 2012) was TCA-precipitated and dried protein pellets were subjected to SDS PAGE as described in Top et al. (2019). The band of EPO was excised and digested simultaneously with trypsin (Promega) and GluC (Thermo Fisher Scientific, Bremen, Germany). Peptides were cleaned as described in Top et al. (2019). MS measurements were performed on RSLCnano system (Dionex LC Packings/Thermo Fisher Scientific, Dreieich, Germany) coupled online to a QExactive Plus instrument (Thermo Fisher Scientific) as described in Top et al. (2019) using 35 % normalized collision energy.

Raw data were processed with Mascot Distiller V2.5.1.0 and database searches on generated peak lists were performed using Mascot Server V2.6.2 and a database containing all *P. patens* protein models V1.6 (Zimmer et al., 2013), the sequence of human EPO (P01588) as well as their reversed sequences used as decoys. Simultaneously, the search was performed against a custom in-house database containing sequences of known MS contaminations such as human Keratin or Trypsin (267 total entries, available on request). Carbamidomethylation (C + 57.021464 Da) was set as fixed modification. Variable modifications were Gln → pyro Glu (N term Q − 17.026549 Da), oxidation (M + 15.994915 Da), Acetylation (N-term + 42.010565 Da), hydroxyproline (P + 15.994915 Da), mono-arabinosylation (P + 148.037173 Da), di-arabinosylation (P + 280.079432 Da) and tri-arabinosylation (P + 412.121691 Da). The peptide mass tolerance was ±5 ppm, and the fragment mass tolerance was ±0.02 Da. Enzyme specificity was set to “none” and a maximum of two missed cleavages was allowed. Results were loaded in Scaffold4 software V4.11.0 using the Legacy Independent Sample Grouping Option and Legacy PeptideProphet Scoring (high mass accuracy).

### 2.8 Identification of secretory proteins and HRGPs

The presence of signal peptides in Physcomitrella proteins was predicted with SignalP 5 (Almagro Armenteros et al., 2019) *via* the library ragp (version 0.3.5.9000; Dragićević et al., 2020) in R (version 4.3.0; R Core Team, 2024). Proteins identified by MS were filtered for the presence of a predicted signal peptide and only those proteins were further considered in the whole study. HRGP classes were assigned based on Liu et al. (2016) and Ma et al. (2017). Physcomitrella V1.6 IDs as used by Liu et al. (2016) were translated into Physcomitrella V3.3 IDs via the PpGML DB (Fernandez-Pozo et al., 2020) and proteins were considered as chimeric AGP if the Physcomitrella V3.3 protein sequence was identical to that of a predicted chimeric AGP from Ma et al. (2017) or if the complete sequence of a predicted chimeric AGP was part of the V3.3 sequence of the measured protein.

### 2.9 Selection of Hyp sites and non-hydroxylated proline sites

Database search results were filtered in Scaffold 4 (or Scaffold 5 in case of the P4H1 KO - P4H6 KO dataset; Proteome Software, Inc., Portland, OR 97219, USA) for a protein and peptide probability > 90% and a minimum number of unique peptides per protein of 1. Only peptides with a Mascot Ion Score > 25 were accepted. Proteins in the results from Mascot searches performed with a database containing Physcomitrella V1.6 protein sequences were translated to the major isoform of Physcomitrella V3.3 proteins using the *P. patens* lookup table downloaded from the PpGML DB (Fernandez-Pozo et al., 2020). If a protein V1.6 ID translated to two V3.3 IDs both were kept and proteins without a V3.3 counterpart were removed. Similarly, peptides that did not fit into the translated V3.3 protein sequence were removed. Translation from V1.6 to V3.3 was performed for the following datasets: host cell proteome (Hoernstein et al., 2018), preliminary measurements of EPO-producing line (174.16), separate measurements of maternal line (174.16) and P4H1 KO line. For all other datasets the Mascot searches were directly performed with a database containing Physcomitrella V3.3 protein sequences.

Results in mzIdentML format were exported from Scaffold and converted to pepXML format with help of the OpenMS software (version 2.7.0, Röst et al., 2016). The probability for the localization of the hydroxylation (+15.99) at a specific proline compared to other prolines as well as methionine and tryptophan that can be hydroxylated as artefacts during electrospray ionization (Silva et al., 2013) was computed with PTMProphet from the Trans-Proteomic Pipeline (TPP v5.2.0; Deutsch et al., 2015). Spectra in which the hydroxylation of proline could not be distinguished from hydroxylation of carbamidomethylated cysteine, which was described to become hydroxylated in Na et al. (2012), as well as histidine, tyrosine and phenylalanine, which are susceptible to oxidation (Berlett and Stadtman, 1997), were not considered further. For peptides that fitted twice in the same protein, the first match was chosen as the Hyp site. If peptides were assigned to two different but very similar proteins one representative protein was selected.

Prolyl-hydroxylation as modification was accepted at a PTMProphet probability > 0.7 at the specific proline. Moreover, a set of prolines that were not measured to be hydroxylated was selected from the MS data of lines without P4H KO, including proline sites with a very low probability of hydroxylation (PTMProphet probability < 0.01).

### 2.10 Computation of the degree of prolyl-hydroxylation

The degree of prolyl-hydroxylation of Hyp-containing peptides was estimated from peptide intensity values of the maternal line in the quantified MS data (data from P4H KO lines was not included). The summed intensities from all forms of a peptide with prolyl-hydroxylation (i.e., all charges and all combinations of post-translational modifications) were divided by the summed intensities of the peptide (prolyl-hydroxylated and not prolyl-hydroxylated version) and the mean over three technical replicates from two datasets was taken.

### 2.11 Hyp amino acid sequence environment

The logo showing the AA frequency in sequence windows of length 15 centred around Hyp sites was created with the standalone version of WebLogo (version 3.7.9, Crooks et al., 2004) setting *probability* as the unit and *none* for composition. The two-sample logo was created with the Two Sample Logo web-based application (Vacic et al., 2006) using as positive samples sequence windows of length 15 centred around Hyp sites and as negative samples sequence windows centred around the selected set of prolines that were not measured to be hydroxylated. The *P* value cut-off was set to the default value of 0.05. In the WebLogo and the two-sample logo the P of the central proline was replaced with an O using GIMP (version 2.10.18).

### 2.12 Prediction of Hyp sites

Prediction of Hyp sites was performed for the major isoforms of all secretory Physcomitrella proteins with predicted signal peptide excluding proteins encoded by plastids or mitochondria. Identification of prolines hydroxylated according to the glycomodule and the extended prolyl-hydroxylation code was done with a Python script. Further, Hyp sites were predicted with the library ragp (0.3.5.9000; Dragićević et al., 2020) in R (version 4.3.0; R Core Team, 2024) using the default probability threshold of 0.224.

### 2.13 Structural analysis

Models of proteins containing validated Hyp sites or prolines from the selected set of non-hydroxylated prolines were downloaded from the AlphaFold Protein Structure Database (Varadi et al., 2022). The relative accessible surface area as well as the six secondary structure elements (3-10 helix, α-helix, π-helix, strand (participates in β ladder), isolated β-bridge, turn (hydrogen bonded), bend) or none of the previous were assigned to each residue with the DSSP module from Biopython (v1.80, Cock et al., 2009).

Models of the six Physcomitrella P4Hs with the bound EPO peptide EAISPPDAASAAPLR were generated with ColabFold (1.3.0; Mirdita et al., 2022). The models were built with AlphaFold2-multimer-v3 (Evans et al., 2022) using no template information. The program was run with default settings and the top-ranking model of P4H1 was relaxed with molecular dynamics. Additionally, models of the six Physcomitrella P4Hs with the bound EPO peptide were generated with AlphaFold2-multimer-v2 (Evans et al., 2022) using no template information and 48 recycles. All further visualizations and analyses were performed in PyMOL (The PyMOL Molecular Graphics System, Version 2.3.0 Schrödinger, LLC): the identification of hydrogen bonds between P4H1 and the substrate peptide, the alignment with the crystal structure from the *Chlamydomonas reinhardtii* P4H1 with a bound peptide substrate downloaded from the PDB (https://www.rcsb.org/; PDB ID: 3GZE chain C; Koski et al., 2009) and the computation of the root-mean-square deviation (RMSD) between the two peptide substrates. The latter was computed with the rms_cur command considering the five C_α_ atoms in the range of two AAs around the proline that becomes hydroxylated.

### 2.14 Protein and peptide abundance

Significant changes in the abundance of Hyp-containing peptides in P4H KO lines (light) compared to the ^15^N labelled maternal line (heavy) were computed with the light/heavy (L/H) intensity ratios for the P4H1 – P4H6 single KO dataset. The L/H ratios of each replicate were log2 transformed and normalized to a median of zero by subtraction of the median. For protein-level analysis the normalization was performed using the median value of the peptides in the replicate. For peptide-level analysis the normalization was performed with the median value of the respective protein. A one sample *t*-test was performed with an expected value of zero using SciPy (1.10.0, Virtanen et al., 2020) and peptides with *P* < 0.05 were accepted. Further, the mean of the normalized log2 transformed L/H ratios of the three replicates was computed and only peptides with a reduced abundance where this value was < 1 were kept. If there was a reduction in the abundance of the unmodified peptide that was comparable to the reduction in the abundance of the Hyp-containing peptide, this peptide was not further considered.

For computation of significant changes in the abundance of proteins in P4H KO lines, the same procedure to compute the *t*-test was applied as for the peptides but with the protein L/H intensity ratios and only proteins present in all three replicates were considered. Afterwards, multiple testing correction was performed using the Benjamini/Hochberg method via the Python module statsmodels (0.13.2, Seabold and Perktold, 2010). Proteins were filtered for an adjusted *P* value < 0.05 and |mean normalized log2 L/H ratio| > 1. Finally, only proteins that had a probability for correct protein identification > 90 % in Scaffold 5 (Proteome Software, Inc., Portland, OR 97219, USA) were selected. Proteins for which the direction of the change in abundance was opposite in the two datasets used for this analysis were removed.

### 2.15 Transcript abundance

In order to quantify the expression levels of P4H genes, RNA was isolated from 100 mg plant material of wild type and the KO lines, respectively. The RNA was first digested with DNAse I. After incubation for 1h at 37 °C in the thermo block, the reaction was stopped by addition of 2 μL EDTA (25 mM) and incubation at 65 °C for 10 min. After DNAse I digest, the RNA was reverse-transcribed to cDNA with TaqMan® Reverse Transcription Kit, using Multiscribe RT in a thermocycler. RT-PCR was performed with appropriate primers (efficiency =2 ± 0.1; calculated by the Abs Quant\2nd Derivate max). The qRT-PCR amplification was performed with SensiMix^TM^ SYBR NO-Rox Kit (Bioline) according to the manufacturer’s recommendations. The gene expression was normalized against the housekeeping genes *EF1α* (Pp3c2_10310V3.1) and *L21* (Pp3c13_2360V3.1) (Wiedemann et al., 2018; Bohlender et al., 2020) and the relative quantification was calculated based on Advance Relative Quantification provided by the LightCycler®480 (software release 1.5, Roche Diagnostics). Finally, the expression of the P4H genes was normalized against the maternal line and statistical analysis of the mean values of the qRT-PCR was performed using the GraphPad Prism software (version 8.0; La Jolla, California, USA) using an ANOVA with Durentt‘s test (*P* < 0.05).

### 2.16 Data analysis and visualisation

For data analysis in Python 3 (v3.8.10, van Rossum and Drake, 2009) the libraries Pandas (v1.3.4, McKinney, 2010; The pandas development team, 2020) and numpy (v1.21.4, Harris et al., 2020) were applied and figures were generated with the libraries Matplotlib (version 3.2.1; Hunter, 2007), Seaborn (version 0.10.1; Waskom, 2021) and pyvenn (version 0.1.3, https://github.com/tctianchi/pyvenn).

## 3. Results

### 3.1 Six Physcomitrella prolyl hydroxylases in four subfamilies

Due to sub- and neofunctionalisation, different enzyme isoforms may have different substrate specificities. Therefore, we evaluated the P4H family by phylogenetic reconstruction and identified six Physcomitrella P4Hs in four distinct clades resembling putative subfamilies in the Viridiplantae (Figure 1, Supplemental Figure S1). Two Physcomitrella P4Hs, namely P4H1 and P4H2, each belong to a distinct subfamily whereas P4H3 and P4H4 as well as P4H5 and P4H6 group pairwise in common clades. All Physcomitrella P4Hs with the exception of P4H4 have direct orthologues in *F. hygrometrica* from the same family of mosses. While we found members of all subfamilies encoded by the mosses *C. plumiforme*, *C. purpureus*, *S. fallax* and *S. magellanicum*, the enigmatic *T. lepidozioides*, sister to all other mosses, only encodes two P4Hs: one orthologue of Physcomitrella P4H1 and one single orthologue of the Physcomitrella P4H5 and P4H6. Homologues of the hornwort *A. angustus* are present in all clades, while we found homologues of the liverwort *M. polymorpha* in all clades but the one containing Physcomitrella P4H1. Of the twelve Arabidopsis P4Hs, AT-P4H-1 is an orthologue of Physcomitrella P4H1, AT-P4H-2, AT-P4H-4, AT-P4H-6 and AT-P4H-7 are co-orthologues of Physcomitrella P4H2, AT-P4H-9 and AT-P4H-13 are co-orthologues of Physcomitrella P4H3 and P4H4, while AT-P4H-3, AT-P4H-5, AT-P4H-8, AT-P4H-10 and AT-P4H-11 are co-orthologues of Physcomitrella P4H5 and P4H6. In addition, we identified eleven P4H proteins within the latest annotation of *N. benthamiana* (Ranawaka et al., 2023), which clustered in congruence with Mócsai et al. (2021) and were labelled accordingly.

**Fig. 1.**
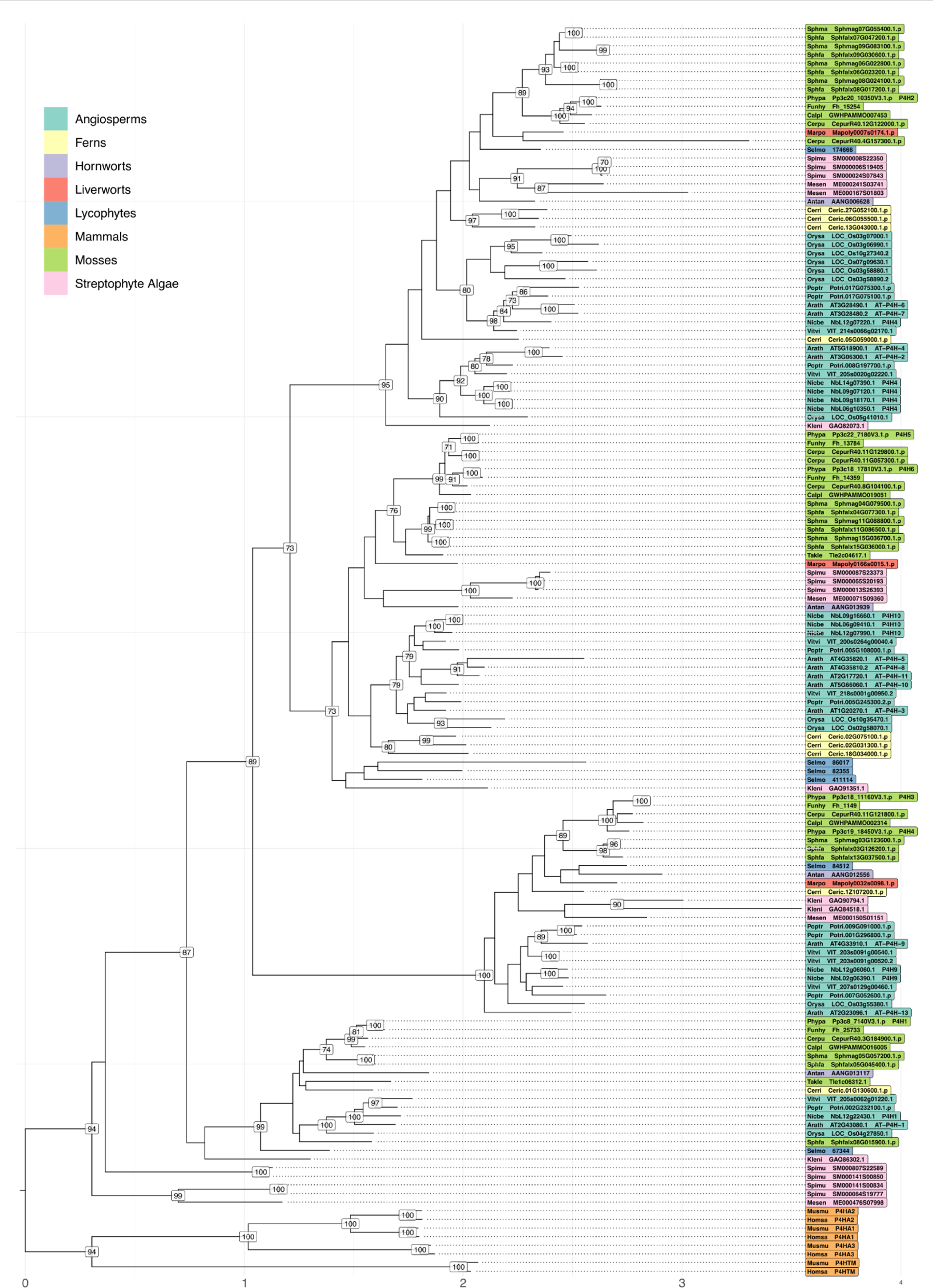
Phylogenetic reconstruction of the plant P4H family. Maximum-likelihood tree of a codon-aware multiple sequence alignment of P4H coding sequences annotated with internal bootstrap support values (>= 70 %) and outgroup-rooted with mammal sequences. Tip labels are coloured by taxonomic units, reference sequences are complemented by their trivial name and species abbreviations follow a five-letter code. Antan: *Anthoceros angustus*; Arath: *Arabidopsis thaliana*; Calpl: *Calohypnum plumiforme*; Cerpu: *Ceratodon purpureus*; Cerri: *Ceratopteris richardii*; Funhy: *Funaria hygrometrica*; Homsa: *Homo sapiens*; Kleni: *Klebsormidium nitens*; Marpo: *Marchantia polymorpha*; Mesen: *Mesotaenium endlicherianum*; Musmu: *Mus musculus*; Nicbe: *Nicotiana benthamiana*; Orysa: *Oryza sativa*; Phypa: *Physcomitrium patens*; Poptr: *Populus trichocarpa*; Selmo: *Selaginella moellendorfii*; Sphfa: *Sphagnum fallax*; Sphma: *Sphagnum magellanicum*; Spimu: *Spirogloea muscicola*; Takle: *Takakia lepidozioides*; Vitvi: *Vitis vinifera*.

### 3.2 Mass spectrometry revealed 73 Hyp sites in 24 secretory proteins

We identified Hyp sites in Physcomitrella proteins using multiple MS/MS measurements. Due to our extraction protocols the samples were enriched for secretory proteins from ER, Golgi apparatus, cell wall and extracellular space. In total, 5139 proteins were measured in the MS data sets. From these, 602 had a predicted signal peptide, that allows them to enter the secretory pathway *via* the ER and get in contact with the P4H enzymes which are localized here (Parsons et al., 2013) and catalyse the prolyl-hydroxylation reaction. The MS data covered 23.3% of all proteins with a predicted signal peptide from the Physcomitrella proteome (Supplemental Figure S2a) but only 6.86% of their proline sites were covered by identified peptides (Supplemental Figure S2b). No signal peptide was predicted for the other 4537 proteins, so it is not certain whether they pass through the secretory compartments. Hyp sites were collected exclusively from proteins with predicted signal peptide. This resulted in 73 validated Hyp sites (PTMProphet probability > 0.7; Supplemental Figure S3) from 26 peptides after trypsin, elastase or thermolysin cleavage belonging to 24 proteins (Supplementary Table T1). Peptide versions with different cleavage sites were not counted additionally. Some of the 26 peptides with validated Hyp sites were also measured without prolyl-hydroxylation. For seven of these the degree of prolyl-hydroxylation was estimated using peptide intensities from the quantified MS data (Supplemental Figure S4). These seven peptides had a varying degree of prolyl-hydroxylation between 0.04 % and 10.83 %. With one exception (AASILLYHIV**O**S**O**ATAADLTDGQTLTTALGK) these sites with low degree of prolyl-hydroxylation were isolated prolines that had no other proline in the neighbourhood of two AAs at each side.

Physcomitrella HRGPs were predicted by Liu et al. (2016) and Ma et al. (2017). According to the classification from these studies, none of the 24 measured Hyp-containing proteins was an extensin but eight of them were chimeric AGPs. These were one laminin G-like AGP (Pp3c1_2420V3.1), two xylogen-like AGPs (Pp3c1_11030V3.1; Pp3c1_31020Vtermh3.1), one phytocyanin-like AGP (Pp3c16_22330V3.1), one fructose-1,6-bisphosphatase-like AGP (Pp3c18_4950V3.1) and three fasciclin-like AGPs (Pp3c4_16840V3.1; Pp3c7_430V3.1; Pp3c21_10620V3.1) comprising together 48 of the measured Hyp sites (Supplementary Table T1). The protein with the highest number of validated Hyp sites (11) was a myosin light-chain kinase of the chimeric xylogen-like AGP family (Pp3c1_11030V3.1). Also, in several of the other chimeric AGPs a high number of Hyps was measured. These were nine Hyps in the chimeric phytocyanin-like AGP (Pp3c16_22330V3.1) as well as seven Hyps each in the chimeric xylogen-like AGP (Pp3c1_31020V3.1) and in one of the three chimeric fasciclin-like AGPs (Pp3c4_16840V3.1). From the 16 Physcomitrella proteins that were no HRGPs, only one Hyp each was identified in 11 proteins, two proteins contained two Hyps, two other proteins contained three Hyps and in one protein four Hyps were detected.

### 3.3 Alanine, threonine, proline, serine and valine are enriched around Hyps

Since the AA in the protein sequence at the position directly before the Hyp (position -1) has a special importance according to established prolyl-hydroxylation motifs for plants (Gomord et al., 2010; Canut et al., 2016), we determined frequencies of each AA which was found immediately before the 73 measured Hyps. The most frequent AAs were in descending order alanine, threonine, proline/hydroxyproline and serine (Figure 2). Valine was slightly more abundant than the remaining AAs that were either counted once or twice before a Hyp (twice: G, K; once: C, I) or not detected at all (D, E, F, H, L, M, N, Q, R, W, Y). Due to the applied filtering criteria, excluding all sites with ambiguous localization of the hydroxyl group on the proline, Hyp sites in proximity to methionine and tryptophan might be underrepresented and to a lesser extent this might also affect the easily oxidable AAs cysteine, histidine, phenylalanine, and tyrosine (Berlett and Stadtmann, 1997).

To investigate which AAs in the protein sequence are present in near and more distant neighbourhood of the Hyps, AA sequence windows of length 15 centred around the Hyp sites were generated. This revealed that the tolerance for the presence of AAs other than proline, alanine, serine, threonine, and valine was clearly the smallest at position -1 directly before the Hyp and increased for positions further apart from the Hyp. Proline/hydroxyproline, alanine and valine (Figure 2a, b) were the AAs that occurred most frequently at several positions of the sequence windows. Indeed, these three AAs as well as threonine and serine were significantly enriched at one or several positions around the Hyps, when using as reference the AA distribution around 2773 prolines sites that were non-hydroxylated (PTMProphet probability for hydroxylation <0.01, sites less than seven amino acids apart from protein termini were not considered, Supplemental Figure S3) in secretory proteins with predicted signal peptide (Figure 2c). In contrast, leucine, by far the most frequent AA in all Physcomitrella proteins with predicted signal peptide (Supplemental Figure S5), as well as asparagine and aspartic acid were among the AAs that were underrepresented at one or several positions around Hyps (Figure 2c). Hereinafter, hydroxylated proline residues within discussed or detected sequence motifs will be depicted as “O” for easier discrimination from non-hydroxylated residues.

**Fig. 2.**
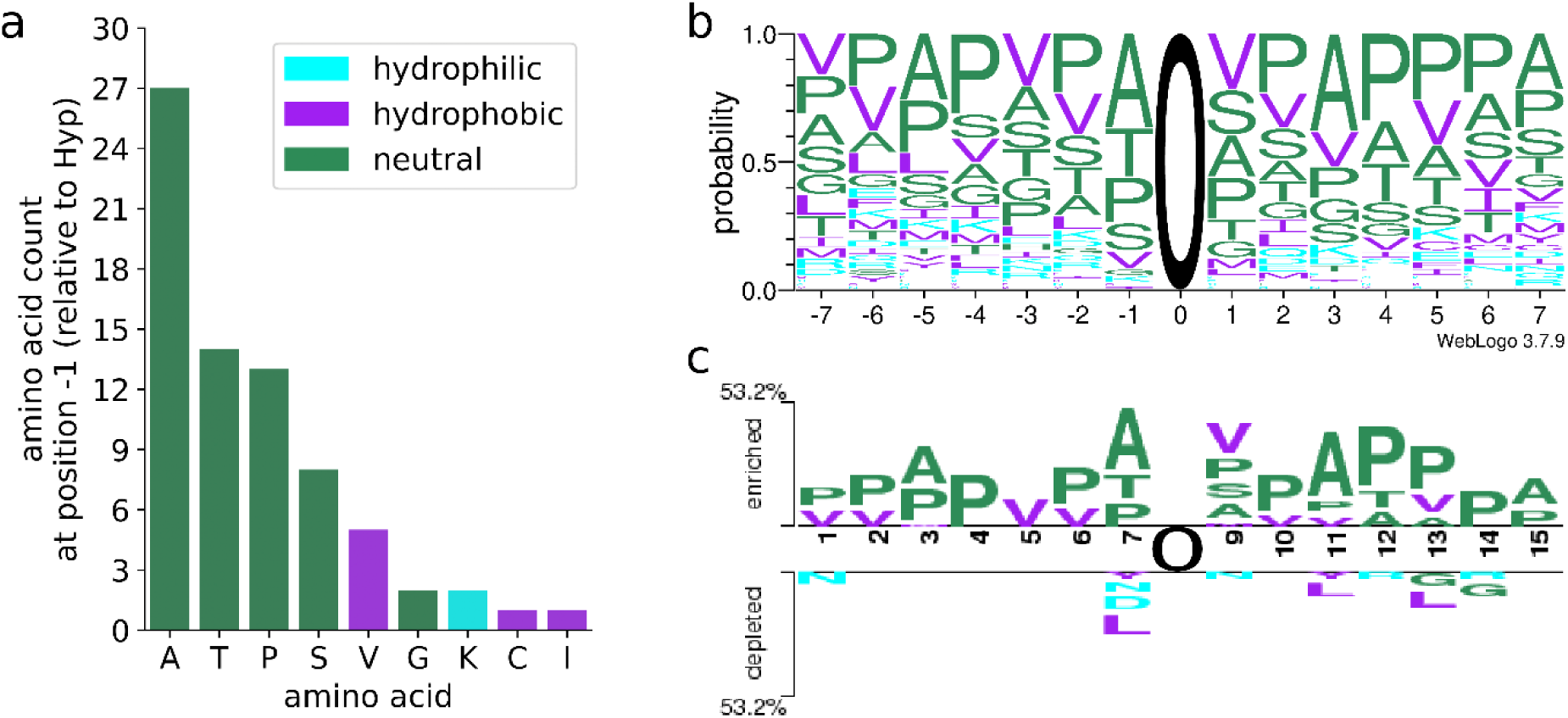
Amino acid distribution around Hyps from Physcomitrella proteins. Depicted is the frequency of different AAs located directly before the identified Hyps (a), the AA distribution in AA sequence windows of length 15 centred at the 73 Hyp sites (b) and a two-sample logo illustrating the AAs that were enriched or depleted around Hyps compared to 2773 proline sites not measured to be hydroxylated (c). In (b) and (c) the central Hyps are depicted as O at position 0. The AAs and corresponding bars are coloured by their properties: green = neutral AAs (A, G, H, P, S, T), purple = hydrophobic AAs (C, F, I, L, M, V, W, Y), cyan = hydrophilic AAs (D, E, K, N, Q, R).

### 3.4 APV is a frequent prolyl-hydroxylation pattern

Of the 73 Hyps, 50 were not surrounded directly by other prolines or Hyps, while the other 23 were part of blocks of two to four amino acids that included combinations of prolines or Hyps. Analysing the combination of the AA before and after a single non-contiguous Hyp, the combination of A**O**V was by far the most frequent and was present in total 15 times (Supplementary Table T2). A**O**V was found in six of the Hyp-containing proteins two of them being no HRGPs and four being chimeric AGPs. In two of the chimeric AGPs (Pp3c4_16840V3.1, Pp3c16_22330V3.1) this combination was part of the repetitive motif (A**O**VV)_3-4_ (Supplemental Figure S6). Other combinations of flanking AAs were A**O**A, T**O**S and V**O**A (four times), as well as A**O**G, A**O**T and T**O**T (three times) that were in chimeric AGPs most often part of the glycomodule motif (Supplemental Figure S6). AA combinations flanking two prolyl-hydroxylation sites were A**OO**M, S**OO**Q, S**OO**S, T**OO**D, T**OO**M, and T**OO**S. Such combinations were found in the longer motif of [A/T]**OO**MGST**OO**S, that was identified three times in one of the chimeric AGPs (Supplemental Figure S6). While each individual proline written as O in the previously mentioned patterns was at least once hydroxylated, in the motif QP**O**K the first proline was never hydroxylated. Blocks with more than two prolyl-hydroxylation sites were only identified in the peptide K**OOOO**S**O**PPK.

### 3.5 Arabinose on two pectinesterases

To identify which Hyps are *O*-glycosylated with arabinoses, modifications of Hyp with one to three arabinosyl residues were searched for in the MS data. In the two tryptic peptides TEGMGIAGT**OO**DDGSSS**O**S**O**STPTCIR (Pp3c5_12660V3.1) and YEAQNSESTVLDTQTLPGGDFSVEAT**O**S**OO**QEATCIR (Pp3c25_760V3.1) from two different cell-wall located pectinesterases, prolines within the glycomodule motif ([Ala ⁄ Ser ⁄ Thr ⁄ Val]-Pro_(1,4)_-X_(-0,10)_-[Ala ⁄ Ser ⁄ Thr ⁄ Val]-Pro_(1,4)_) were prolyl-hydroxylated and *O*-glycosylated with arabinose residues (Supplementary Figures S7, S8). Most frequently, two arabinose residues were present in the peptides, however, in the first peptide up to five arabinoses were found.

In order to obtain data about peptides containing Hyp sites *O*-glycosylated with arabinogalactans, that normally prevent the detection of these peptides by MS, two approaches for deglycosylation were applied: One was employing a Physcomitrella mutant with a double KO of galactosyltransferases (Δ*galt*2/3) which might be responsible for *O*-glycosylation (Parsons et al., 2012), and the second was chemical deglycosylation and treatment with three different proteases (trypsin, elastase, thermolysin) to increase the sequence coverage of the identified proteins. MS^2^ spectra from eight of the 26 Hyp-containing peptides were exclusively obtained from the deglycosylated datasets (chemical deglycosylation: six peptides, galactosyltransferase KO: two peptides; Supplementary Table T1). All peptides contained at least one Hyp within a glycomodule and some contained a long glycomodule spanning many Hyps (e.g., LVA**O**V**O**A**O**VVKA**O**A**O**A**O**VIKA**O**T**O**G**O**A), making these suitable candidates for *O*-glycosylation.

### 3.6 Combination of three tools predicts about 95% of the Hyp sites

We searched for possible prolyl-hydroxylation sites in all 1920 secretory Physcomitrella proteins (major isoforms) with a predicted signal peptide (no organelles) using three methods appropriate for plant-like prolyl-hydroxylation: the glycomodule ([Ala ⁄ Ser ⁄ Thr ⁄ Val]-Pro_(1,4)_-X_(-0,10)_-[Ala ⁄ Ser ⁄ Thr ⁄ Val]-Pro_(1,4)_), the extended prolyl-hydroxylation code and the R package ragp using default settings (Gomord et al., 2010; Canut et al., 2016; Dragićević et al., 2020). This resulted in 8249 predicted Hyp sites from the glycomodule, 16,546 from the extended prolyl-hydroxylation code and 8414 from ragp. A high number of 4095 proline sites was predicted to be hydroxylated in accordance with all three methods (Figure 3).

Subsequently, the data from the measured secretory proteins with a predicted signal peptide were used to check whether the three methods correctly predict the 73 MS-verified Hyp sites and whether they predict no hydroxylation of the 2828 prolines that we identified as non-hydroxylated. The prediction performance was assessed using the balanced accuracy score (0=no correct prediction; 1=all predictions correct). From the three methods, the ragp tool and the glycomodule performed best and had a comparable performance with 56 (76.71 %) and 55 (75.34 %) correctly predicted Hyps (Figure 3a). The ragp tool, however, predicted a possible hydroxylation for 181 additional proline sites that were not hydroxylated, while 237 were predicted by the glycomodule (Figure 3b), leading to a slightly higher balanced accuracy of the ragp tool compared to the glycomodule (0.85 and 0.83, respectively). The extended prolyl-hydroxylation code correctly predicted 54 of the Hyps (73.97 %; Figure 3a) and a possible hydroxylation for 764 (27.02 %) further proline sites that were measured to be non-hydroxylated (Figure 3b), resulting in a balanced accuracy score of 0.73. Only four identified Hyp sites were not predicted by any of the three methods (Figure 3a, Supplemental Figure S9). One of them (YYPPFK**O**ELVK) was in close proximity to a proline-proline sequence segment, while another one (GHEG**<u>O</u>**SSVYT**O**SSDTEPFNFHDPR, underlined) was the first Hyp in a Gly-Hyp-Xaa_4_-Thr-Hyp motif. The third Hyp site (WNSNIVVVGVDDI**O**LR) was from a chimeric laminin G-like AGP but it was at an isolated proline more than 140 AAs distant from the proline-rich region of the protein (Supplemental Figure S6). The fourth was 10 AAs behind a short proline-rich segment of the protein (…SPNPPNPGPTPPSPPPPEVICDKWR<u>TC**O**AENTCCCTFPVGK</u>…, Hyp-containing peptide is underlined).

Next, we tested to what extent a combination of two or all three methods could improve the prediction performance either by a higher number of correctly predicted Hyps or an increased balanced accuracy score. When considering a proline as being hydroxylated based on any of the three methods, the vast majority of the measured Hyps (69 out of 73 = 94.52 %) were predicted with a balanced accuracy of 0.82. When a proline was only considered to be hydroxylated after prediction by all three methods, far fewer of the measured Hyps (42) were predicted, but compared to the other methods also the smallest number of prolines was incorrectly predicted as Hyps (58), making this combination the most precise. Considering a proline as being hydroxylated after prediction by either ragp or the glycomodule, the two methods with the highest accuracy, a balanced accuracy of 0.86 was achieved – better than that of ragp or the glycomodule alone – and more of the experimentally verified Hyps were predicted (62).

**Fig. 3.**
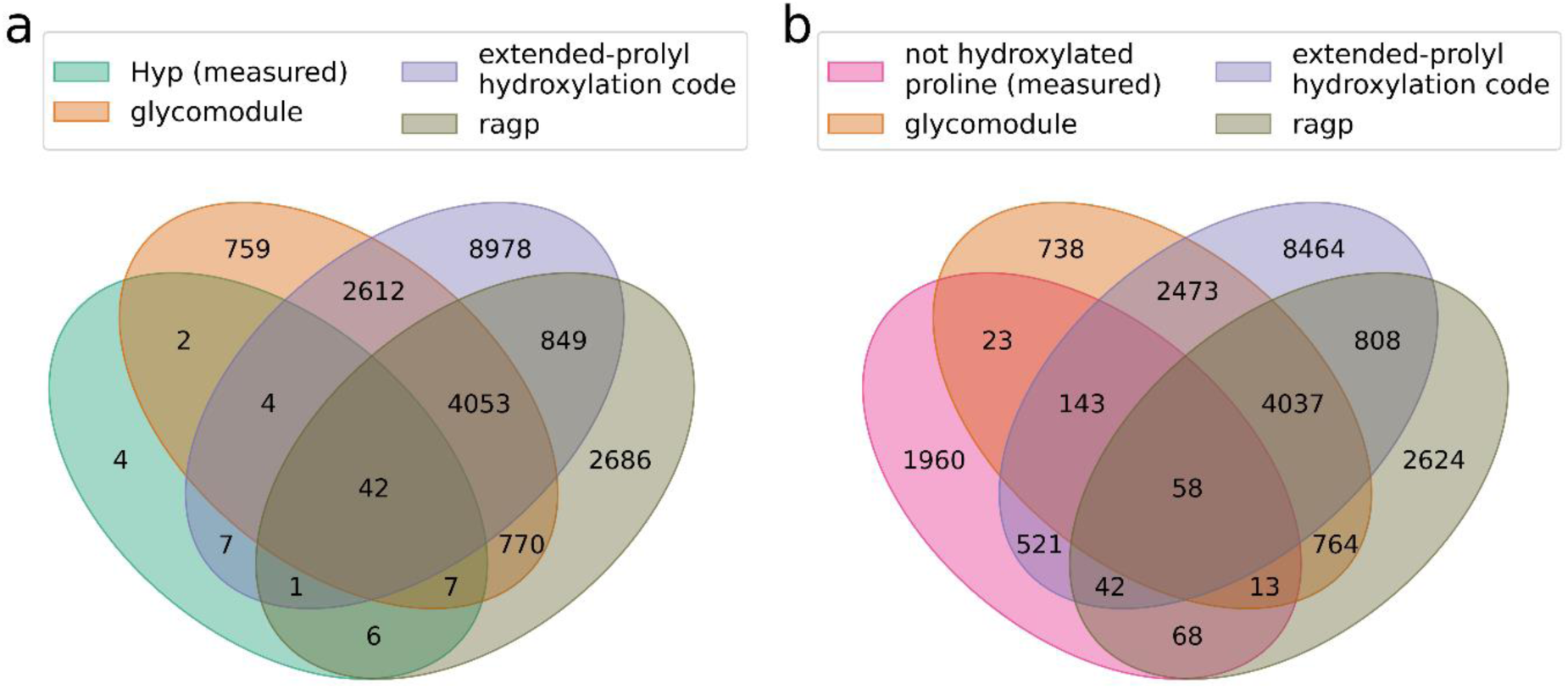
Overlap between Hyps predicted by three methods and measured hydroxylation status of prolines. Hyp sites in all secretory Physcomitrella proteins with a signal peptide (SignalP 5) were predicted using the ragp tool, the glycomodule and the extended prolyl-hydroxylation code. Depicted is the overlap between predicted Hyps sites with measured Hyp sites (a) and proline sites not measured to be hydroxylated (b).

### 3.7 Hyp sites predominantly in disordered regions

To analyse if specific structural features favour prolyl-hydroxylation, 443 protein structure models of secretory proteins with predicted signal peptide, containing the 73 measured Hyps and 2828 proline sites that were non-hydroxylated, were downloaded from the AlphaFold Protein Structure Database (Varadi et al., 2022). The pLDDT confidence score of the model (0=minimum quality, 100=maximum quality) was for most of the non-hydroxylated proline residues higher than 90 (2026 out of 2828 sites), indicating a high quality of the local model structure, whereas it was below 50 for 42 of the 73 Hyp sites (Supplemental Figure S10). Low pLDDT scores can be an indication for disorder (Tunyasuvunakool et al., 2021).

The DSSP tool was used to determine where the protein structure model was folded into any of the seven secondary structure elements (3-10 helix, α-helix, π-helix, strand (participates in β ladder), isolated β-bridge, turn (hydrogen bonded), bend). Additionally, the relative accessible surface area (0=completely buried within protein structure, 1=fully exposed to solvent) of each residue was computed. The Hyps were located mostly in well accessible protein regions with a median relative accessible surface area of 0.86, whereas non-hydroxylated prolines were often less accessible, having a median relative accessible surface area of 0.37 (Supplemental Figure S10). Considering the secondary structure of the protein, Hyps were only present in four of the defined secondary structure elements (in bends, 3-10 helices, strands and turns; Supplemental Figure S10), but most frequently both, Hyps and non-hydroxylated prolines, were located in regions of the protein models where none of the seven secondary structure elements were assigned (Supplemental Figure S10). With 87.67 % the proportion of Hyps in regions without assigned secondary structure was much higher than for the non-hydroxylated prolines with 43.71 %. With five exceptions, the regions without assigned secondary structure containing the Hyp sites were spanning more than 10 and up to 296 AAs (Figure 4c, Supplemental Figure S11, Supplemental Figure S12), while they were shorter than 10 AAs for the vast majority of the non-hydroxylated proline sites (1139 out of 1236 sites without secondary structure; e.g. in Figure 4a and Supplemental Figure S11).

In the chimeric AGPs the long, disordered regions containing the Hyps were mostly rich in prolines and glycomodules while non-hydroxylated prolines were found predominantly in the structured domain of the chimeric AGP (e.g., Figure 4c, Supplemental Figure S11). An exception was for example a Hyp in a chimeric AGP, a xyloglucan endo-transglycosylase (WNSNIVVVGVDDI**O**LR, Pp3c1_2420V3.1) where according to the NCBI Conserved Domain Search webtool (CD-Search; Marchler-Bauer and Bryant, 2004) the single Hyp site is located in the active site of the protein. In the proteins that were no HRGPs, Hyps were for example found in short glycomodules in the N-terminal disordered regions from two pectinesterases (Pp3c25_760V3.1 and Pp3c5_12660V3.1, Figure 4b, Supplemental Figure S12) where the Hyps were *O*-glycosylated with arabinoses.

**Fig. 4.**
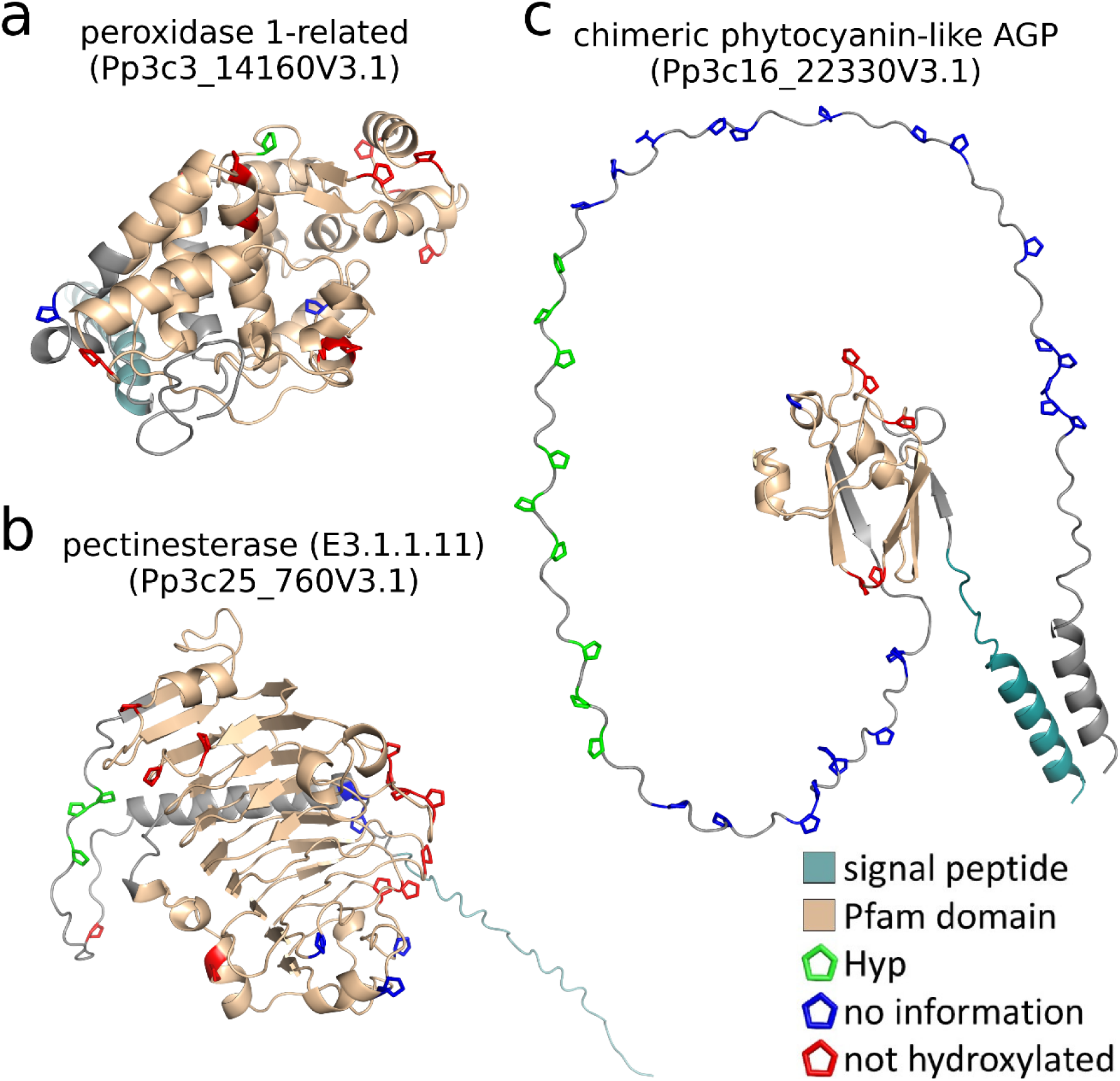
Three-dimensional structures of Hyp-containing proteins. Measured Hyp sites (green) and non-hydroxylated proline sites (red) are highlighted in two exemplary structures from secretory proteins not predicted to be HRGPs (Pp3c3_14160V3.1 in a and Pp3c25_760V3.1 in b) as well as a chimeric phytocyanin-like AGP (Pp3c16_22330V3.1 in c). All remaining prolines are coloured blue. For these, no certain information about their hydroxylation status could be gained from the MS data. The Pfam domains for peroxidase (PF00141 in a), pectinesterase (PF01095 in b) and the plastocyanin-like domain (PF02298 in c), respectively, as given by Phytozome (v13; Goodstein et al., 2012), are coloured in light brown.

### 3.8 Physcomitrella-produced EPO with plant-specific *O*-glycans

We did not only identify Hyps in native Physcomitrella proteins, but also in Physcomitrella-produced recombinant human EPO. In accordance with Parsons et al. (2013), we found prolines of the EPO peptide EAISPPDAASAAPLR to be hydroxylated. In addition, post-translational modifications of the peptide EAISPPDAASAAPLR included not only prolyl-hydroxylation but also plant-specific *O*-glycosylation of Hyps with arabinose chains on the Ser-Pro-Pro motif (Supplemental Figure S13) as well as, in some occasions, an additional glycosylation of the first serine with a single hexose, if neighbouring prolines were hydroxylated and *O*-glycosylated. From the 1013 spectra of the respective peptide, in 18.86% (191 spectra) the peptide was glycosylated, mostly with more than one arabinose, in 48.17% (488 spectra) the peptide was just hydroxylated (one, two or three Hyps; no arabinose) and in 32.97% (334 spectra) the peptide was unmodified (Figure 5). All the hydroxylated prolines in the EPO peptide fit the glycomodule, but only the second and third proline were predicted to be hydroxylated by the ragp tool with probabilities of 0.33 and 0.26, respectively. Two further Hyp sites in EPO were predicted by the ragp tool at AA sequence positions 29 and 30 in an Ala-Pro-Pro motif. These prolines were not covered in the MS data, but Parsons et al. (2013) reported these as non-hydroxylated.

**Fig. 5.**
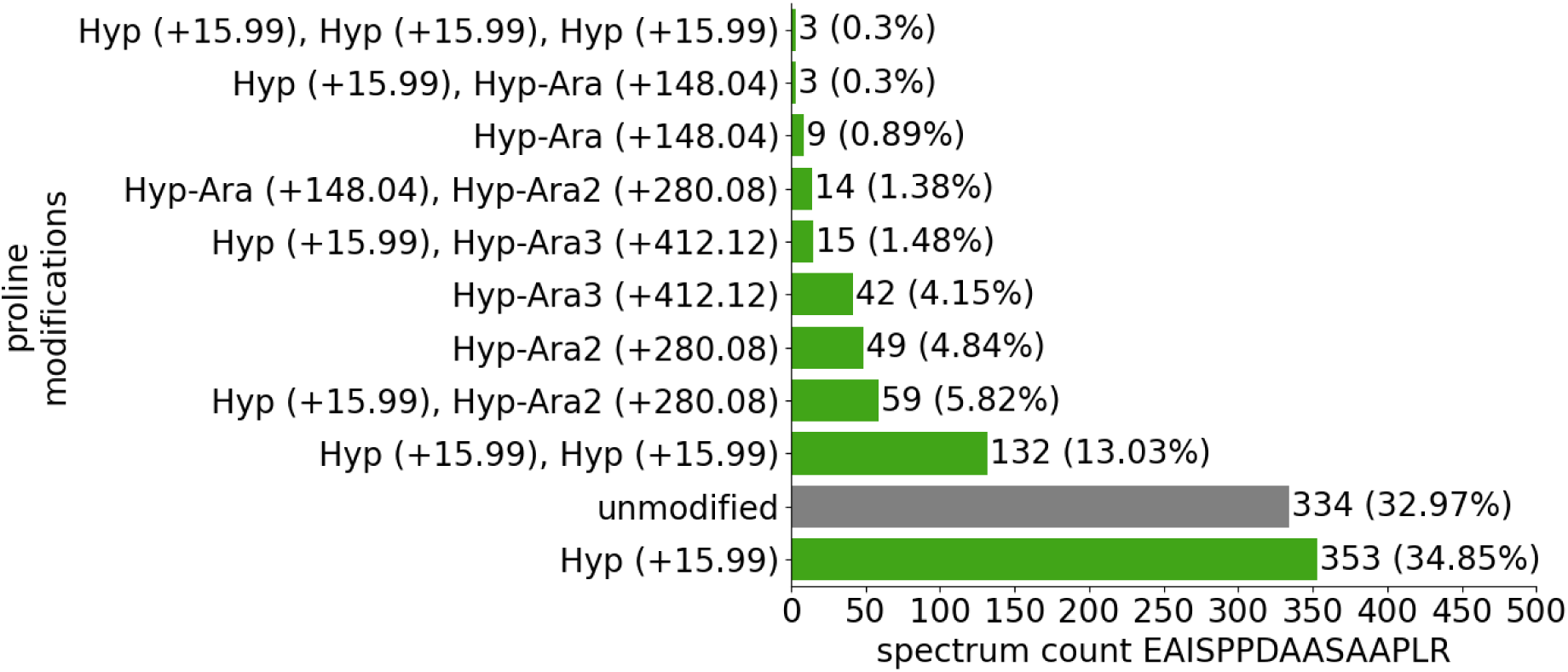
Modified versions of the EAISPPDAASAAPLR peptide from recombinant EPO. Depicted are the number of spectra of the unmodified, prolyl-hydroxylated and the arabinosylated versions of the peptide, respectively. The spectra are counted over several MS measurements and replicates.

### 3.9 Multiple effects of single P4H knockouts

To analyse the effects of the six Physcomitrella P4Hs more deeply, we employed KO mutants to examine prolyl-hydroxylation and abundance of moss proteins. Samples enriched with secretory proteins (from ER, Golgi, cell wall and extracellular space) were obtained using two different protocols. The samples from the EPO-producing maternal line (174.16) were labelled with ^15^N (heavy) and mixed with the single KO lines of each of the six P4Hs (light), respectively, prior to MS measurements and quantification. Significant changes in protein abundance were determined over the light/heavy ratios (= intensity in P4H KO line / intensity in maternal line) in three replicates *via* a *t*-test (p adjusted < 0.05, |log2 light/heavy ratio|>1) and filtered for secretory proteins with a signal peptide. Combining results from both protocols, the P4H6 KO dataset contained the highest number of proteins with altered abundance compared to the maternal line (52 proteins), whereas the smallest number of proteins with altered abundance were 12 proteins in the P4H4 KO dataset (Supplemental Figure S14). Considering the direction of the change, many more proteins had an increased abundance in the P4H1 KO and P4H2 KO dataset while the number of proteins with reduced and increased abundance, respectively, was balanced in the other P4H KO datasets. Some proteins had an altered abundance only in one specific P4H KO dataset while it differed for others in multiple of the P4H KO datasets (Supplemental Figure S14). Only one protein, a subtilisin-like protease (Pp3c11_4360V3.1), was significantly altered (increased) in all P4H KO datasets. A galactose oxidase (Pp3c10_8570V3.1) and a polygalacturonase (Pp3c21_6170V3.1) had an increased abundance in five of the P4H KO datasets, with the latter being strongly increased with a maximal log2 light/heavy ratio of 22.87 in the P4H6 KO. Moreover, the abundance of four chimeric AGPs was increased in one or several of the P4H KO datasets (Pp3c1_2420V3.1, Pp3c4_16840V3.1, Pp3c16_22330V3.1, Pp3c26_5590V3.1, Pp3c4_3520V3.1), and the abundance of one (Pp3c4_3520V3.1) was decreased (Supplementary Table T3).

Additionally, we used these datasets to search for peptides from secretory Physcomitrella proteins with a predicted signal peptide where the abundance of peptide versions with prolyl-hydroxylation was significantly reduced in the P4H KO mutants compared to the maternal line (*P* < 0.05, log2(light/heavy intensity ratio)<1), while the abundance of the corresponding protein and, if present, that of the unmodified peptide were not significantly reduced. In these datasets eight of the previously collected peptides with validated Hyp sites were measured in more than one replicate and hence appropriate for statistical evaluation. We identified a single peptide, GANYAITFCPTVT**O**VAK from a thaumatin family protein (Pp3c16_17280V3.1) in the P4H5 KO line with a reduction in its abundance by a log2 light/heavy ratio of -9.57 (Supplementary Table T4).

Considering the EPO peptide EAISPPDAASAAPLR (or ALGAQKEAISPPDAASAAPLR) a clear trend for reduction in the abundance of its prolyl-hydroxylated form was visible in the P4H1 KO line (Supplemental Figure S15). In all cases the reduction of Hyp-containing or arabinosylated peptides was more than 90%, e.g., the abundance of the peptide with hydroxylation on the second proline (EAISP**O**DAASAAPLR) was reduced by a log2 light/heavy ratio of -4.09 (Supplementary Table T4). In none of the other P4H KO lines the mentioned filtering criteria for significant reduction in the abundance of this peptide were fulfilled and no major changes in prolyl-hydroxylation of the peptide were observable (Supplemental Figure S15). These findings further support the major role of P4H1 in the hydroxylation of EPO in Physcomitrella.

To study if the KO of a single P4H influences the expression of the five other Physcomitrella *p4h* genes, resulting in a putative complementation of the respective KO phenotypes, transcript abundances of each *p4h* gene were determined in the maternal line (174.16) and the six P4H single KO lines. Significance in the changes of transcript abundance was computed with ANOVA and Durentt‘s test (*P* < 0.05). The KOs of the *p4h1*, *p4h2* and *p4h4* genes, respectively, did not significantly change the expression of any of the remaining *p4h* genes. However, our data showed an increase of *p4h1* transcript abundance in the *p4h3* KO, whereas in the *p4h5* KO both *p4h1* and *p4h2* had an increased abundance, while the *p4h6* KO led to an increased transcript abundance of *p4h5* (Supplemental Figure S16).

### 3.10 Modelling suggests peptide interactions in the active site for P4H1

Since AlphaFold-Multimer can predict the interaction between protein-peptide complexes (Johansson-Åkhe and Wallner, 2022), we modelled the interaction between P4H1 and its target sequence, the EPO peptide EAISPPDAASAAPLR, with AlphaFold2-multimer-v3 and no template information. The top-ranking model structure was superposed with the experimentally solved crystal structure of *Chlamydomonas reinhardtii* (Chlamydomonas) P4H1, that has a PSPS**P**SPS peptide bound in its active site (PDB ID 3GZE chain C; Koski et al., 2009), which adopts a left-handed (poly)L-proline type II helix. The third proline in this peptide (bold) is located at the catalytic active position within the active site of Chlamydomonas P4H1, where the prolyl-hydroxylation reaction takes place and is buried under two loops of the enzyme. In the superimposed model of Physcomitrella P4H1, the second proline from the EPO peptide (EAISP**P**DAASAAPLR) is located at this position (marked with an arrow in Figure 6a, b). The root-mean-square deviation (RMSD) of the C_α_ atoms from the five residues of the two substrate peptides centred within the active site (PS<u>PS**P**SP</u>S and EAI<u>SP**P**DA</u>ASAAPLR) is 0.4 Å, indicating a highly similar fold between the backbones of two substrate peptides within the active site. The EPO peptide interacts over eight hydrogen bonds with residues of the P4H1 protein (Figure 6b). Five of these are also present in the interaction between Chlamydomonas P4H1 and its peptide. Two are located between the central proline that becomes hydroxylated and the ARG197 from P4H1 (corresponding to ARG161 in Chlamydomonas P4H1), two are located between residues of the peptide with VAL116 and one with TYR178 of P4H1 (corresponding to VAL80 and TYR140 in Chlamydomonas P4H1).

To test if the EPO peptide is a possible substrate for any of the other five Physcomitrella P4Hs, those were modelled with the EPO peptide using versions 2 and 3 of AlphaFold2-Multimer. While the peptide fits into the P4H1 models computed with both versions, it was not modelled into any of the other Physcomitrella P4H proteins with AlphaFold2-Multimer-v2. In contrast, AlphaFold2-Multimer-v3 computed models for P4H2, P4H5 and P4H6 with either the first (P4H2, P4H5) or the second proline (P4H2, P4H5, P4H6) of the peptide in the catalytic active position. For P4H3 and P4H4 a part of the peptide was modelled at the active site, but other AAs than proline were placed at the catalytic active position (Supplemental Figure S17). Thus, also AlphaFold2-Multimer-v3 computed P4H-EPO interactions less favourable for the other five Physcomitrella P4Hs than for P4H1. We consider this as further indication for different substrate specificities of the moss P4Hs, and as support of our experimental findings.

**Fig. 6.**
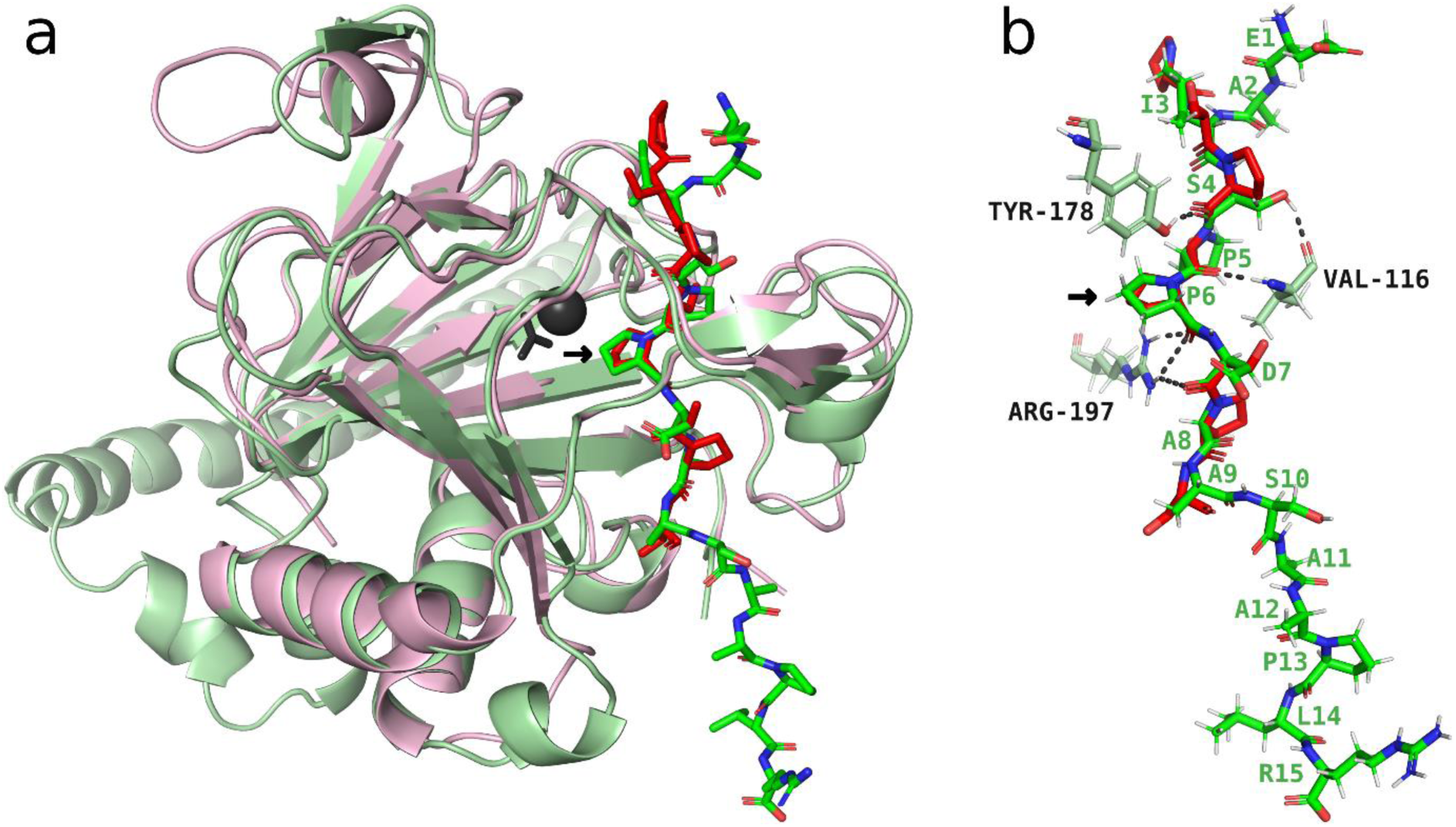
Interaction between P4H1 and EPO peptide EAISPPDAASAAPLR modelled with AlphaFold2-multimer-v3 without template information. (a) The top-ranking model (light green) with the bound peptide EAISPPDAASAAPLR (green) superimposed on the experimentally solved crystal structure of *Chlamydomonas reinhardtii* P4H1 (light red) having a bound (Pro-Ser)_4_ peptide (red) in its active site (PDB ID 3GZE chain C; Koski et al., 2009). (b) Conformation of the two substrate peptides in the superposed structures with hydrogen bonds (dashed lines) between the EPO peptide and P4H1 to the residues VAL80, TYR140 and ARG197.

## 4. Discussion

Plants are gaining increasing importance for the production of valuable compounds, such as pharmaceuticals. While most production hosts are vascular plants, such as *Nicotiana benthamiana* (Nicotiana) and *Daucus carota* (carrot), the non-vascular moss Physcomitrella has a proven track-record for the production of pharmaceuticals and bioactive ingredients (Decker and Reski, 2020; Verdú-Navarro et al., 2023; Munoz et al., 2024). In our attempts to constantly optimize this moss for molecular farming approaches (Reski et al., 2015; Ruiz-Molina et al., 2023), we concentrate on gene expression (e.g., Top et al., 2021; Niederau et al., 2024), bioproduction (e.g., Ruiz-Molina et al., 2022), and glycoengineering (e.g., Decker and Reski, 2012; Bohlender et al., 2022). In the latter field, plant-typical glyco-structures have been abolished (Parsons et al., 2012), and stable *in-vivo* protein sialylation has been achieved (Bohlender et al., 2020). In contrast to the well-studied *N*-glycosylation of recombinant proteins, *O*-glycosylation is still underexplored in plant production hosts although it might deteriorate product quality. While mosses, such as Physcomitrella, and vascular plants, such as Nicotiana, share similar *N*-glycosylation patterns (Koprivova et al., 2003), they may differ in their *O*-glycosylation pattern. A gene responsible for prolyl-hydroxylation of recombinant EPO from Physcomitrella has been identified (Parsons et al., 2013), but a reliable bioinformatic tool to predict this protein modification was not yet available.

Based on genome information we made a phylogenetic reconstruction of plant P4Hs, the enzymes responsible for prolyl hydroxylation. The Physcomitrella genome encodes six P4Hs in four subfamilies, indicating neofunctionalisation during evolution. P4H1 is the only homologue of Physcomitrella within its subfamily and clusters with P4H1 from Arabidopsis. Arabidopsis P4H1 hydroxylates poly-proline and proline-rich motifs in plant proteins, and motifs of the human hypoxia-inducible factor and collagen-like peptides (Hieta and Myllyharju, 2002) that are substrates for mammalian P4Hs (Gorres and Raines, 2010). Arabidopsis P4H2 (Tiainen et al., 2005) clusters with Physcomitrella P4H2 and preferentially hydroxylates substrates with three neighbouring prolines. Arabidopsis P4H5 clusters with Physcomitrella P4H5 and P4H6 and hydroxylates all except the last proline in Ser-(Pro)_4_ extensin motifs (Velasquez et al., 2015). Another P4H from this cluster is Arabidopsis P4H3 that plays a role in the response to low oxygen (Konkina et al., 2021).

To gain a better understanding of the favoured targets for prolyl-hydroxylation by the six Physcomitrella P4Hs, we collected Hyp sites from a set of MS/MS measurements. Since the P4H enzymes are located in secretory compartments, we focused on secretory proteins with predicted signal peptide. With 48 out of the 73 identified Hyps, the majority originated from chimeric arabinogalactan proteins (AGPs) and were mostly located in the disordered part of the protein. AGPs are defined by domains with a high content of proline, alanine, serine and threonine (PAST), and non-contiguous prolines in repetitive motifs preceded by alanine, serine and threonine are frequently hydroxylated (Leszczuk et al., 2023). Accordingly, in our study these AAs and additionally valine dominated over a window of 15 AAs centred around the Hyps.

In agreement with MS measurements of 114 Hyps from 62 glycoproteins in rice (Liang et al., 2020), we found alanine most often to precede Hyps. Valine was the second most frequent AA before Hyp in rice. In contrast, it was the fifth most abundant in our data. Leucine, that was significantly depleted before the Hyps compared to non-hydroxylated prolines in Physcomitrella, was the third most frequent AA preceding Hyps in rice. Other AAs such as aspartic acid, glutamic acid and glutamine were not identified in our MS data, although peptides with prolyl-hydroxylation after these AAs occur in vascular plants, e.g., in *Zea mays* and *Echinacea purpurea* (Canut et al., 2016).

Most Hyps were non-contiguous and thus not directly surrounded by other prolines. Combinations of A**O**V, A**O**A, T**O**S, V**O**A, A**O**G, A**O**T and T**O**T were most frequent before and after a non-contiguous Hyp. These combinations were found particularly often in chimeric AGPs and lay within long glycomodule motifs, spanning multiple Hyps. Most of these peptides were from data where deglycosylation was performed to allow identification of peptides that were likely *O*-glycosylated with large arabinogalactan chains, that cannot be identified by MS. Taken together, these peptides are suitable candidates for *O*-glycosylation in Physcomitrella. Interestingly, two peptides that contain several prolines in the glycomodule with arabinose residues were no HRGPs but two pectinesterases.

Some of the Physcomitrella mutants used in this study were producing recombinant human EPO. In the EPO peptide EAISPPDAASAAPLR we found not only the previously reported prolyl-hydroxylation of the first two prolines (Parsons et al., 2013), but in rare cases also hydroxylation of the third proline as well as *O-*glycosylation with up to three arabinose residues. In agreement with the Hyp contiguity hypothesis that predicts addition of arabinogalactans to single non-contiguous Hyps and arabinose chains to neighbouring contiguous Hyps (Kieliszewski and Lamport, 1994), the arabinose chains were assigned to the segment with two neighbouring contiguous prolines. Further, in a small fraction of the peptides, the serine of the Ser-Pro-Pro motif was glycosylated with a hexose, resembling the *O*-glycosylation pattern in extensins where the hexose attached to the serine is a galactose (Saito et al., 2014).

Three methods to predict Hyp sites in plants developed with data from various plant species, but not mosses, were used to predict the hydroxylation status of the prolines in secretory Physcomitrella proteins: the glycomodule, the extended prolyl-hydroxylation code and the ML-based prediction tool ragp. More than 4000 candidate sites for prolyl-hydroxylation were predicted in accordance with all three methods, indicating that our MS data represents only a small fraction of the total hydroxylation pattern in Physcomitrella. Comparing the predictions by ragp and the glycomodule with MS data, both methods performed comparably well. A combination of the predicted Hyps by ragp and the glycomodule yielded an even higher number of correctly predicted Hyps with a better accuracy, making these methods well suited for the prediction of prolyl-hydroxylation in Physcomitrella. All except four of the identified Hyps were correctly predicted by at least one method. For two of these peptides the degree of hydroxylation was determined, and these peptides were only very rarely prolyl-hydroxylated. The peptide EVQLINIINAPLQGFK contained only in 0.09 % a Hyp and WNSNIVVVGVDDIPLR was in 2.11 % prolyl-hydroxylated, indicating that these two peptides are no preferred targets of P4Hs in moss.

Since different P4Hs can act on the same peptide and to some extent on the same prolines, but with diverging preferences (Mócsai et al., 2021), we investigated the effect of the knockout (KO) of a single *p4h* on the expression of the remaining *p4h* genes. Mostly the expression level of the five remaining *p4h* genes was not significantly altered by the KO of a single P4H, but we found hints for a possible compensation in the single KO lines of P4H3, P4H5 and P4H6 by an upregulation of one or two other *p4h* genes. However, functional compensation apart from transcriptional upregulation is also considerable since all *p4h* genes are expressed in protonema under standard growth conditions. A possible rebalancing effect by the remaining P4Hs was also reported after a quadruple KO of the Nicotiana P4H4 subset, where the KO led to a reduced abundance of the unmodified version of the hinge region from a recombinant IgA1 antibody and to an increased fraction of peptides *O*-glycosylated with pentoses (Uetz et al., 2022). In our data, the abundance of the prolyl-hydroxylated form of the peptide GANYAITFCPTVTPVAK was strongly reduced in the P4H5 KO line, indicating that in this case loss of P4H5 is hardly compensated. In addition, we confirmed the findings of Parsons et al. (2013) that P4H1 plays the major role in the prolyl-hydroxylation of recombinant EPO.

The KO of single P4Hs not only affected prolyl-hydroxylation but also resulted in altered abundance of secretory proteins. Among these was a polygalacturonase with strongly increased abundance in several P4H KO datasets. Furthermore, five HRGPs showed increased abundance in one or several P4H KO mutants, except for one whose abundance was decreased in the P4H6 KO. Increased abundances of HRGPs in P4H KO mutants might be partially caused by a reduction in the *O*-glycosylation with large arabinogalactan trees of some peptides, that prevent sufficient solubilization by our extraction method. This is an indication that the deleted P4H contributes considerably to the prolyl-hydroxylation of the respective HRGP. An effect of a P4H KO on cell wall protein expression was also observed in Arabidopsis where *AGP12* was downregulated in a P4H3 mutant, indicating that the presence of P4Hs is linked with transcriptional regulation of AGPs (Konkina et al., 2021).

By combining 443 AlphaFold structural protein models with our MS data of peptides with nearly 3000 proline sites, we identified Hyps predominantly on accessible protein surfaces in disordered regions of the protein. AlphaFold-Multimer models of Physcomitrella P4Hs with an EPO peptide as substrate suggested a highly accurate structure and identified relevant amino acids in the active centre of P4H1 that form H-bonds with the peptide substrate. In contrast, these models were far less clear about substrate binding for the other Physcomitrella P4Hs, further supporting the differential prolyl hydroxylation of the six moss enzymes.

## Conclusions

Here, we provide a comprehensive analysis of prolyl-hydroxylation in the secretome of the moss Physcomitrella, an established production host for pharmaceuticals. We confirmed that general rules for prolyl-hydroxylation derived from vascular plants also apply to the majority of Hyp sites in moss. Nevertheless, some Hyp sites had an amino acid environment diverging from common motifs and were not predictable by existing methods, demonstrating specific differences in the prolyl-hydroxylation capacity between Physcomitrella and vascular plants. The substrate specificity of the different P4H enzymes is still scarcely known in most plant species. While we demonstrate that some proline sites are mainly hydroxylated by a single P4H, there is also evidence for a compensation of such a P4H KO by increased expression of the other *p4h* genes. To what extent an interplay between the P4H enzymes, such as hetero-dimerization, as observed in Arabidopsis (Velasquez et al., 2015), or overlapping substrate specificities, as reported for Nicotiana (Mócsai et al., 2021), play a role for hydroxylation in Physcomitrella has to be determined. An exact understanding of the conditions for the hydroxylation of a proline by one or several P4Hs will facilitate the modification of prolyl-hydroxylation and *O*-glycosylation and can enhance quality and human compatibility of plant-produced pharmaceuticals.

## Author contributions

C.R. performed research, analysed data and wrote the manuscript. S.N.W.H. and N.v.G. performed research, analysed data and helped writing the manuscript. A.W.G., R.P.S and A.B. performed research. L.L.B. and J.P. performed research, analysed data and helped writing the manuscript. E.L.D. and R.R. supervised research, analysed data, acquired funding and wrote the manuscript. All authors read and approved the final version of the manuscript.

## Acknowledgements

This study was supported by the Ministry of Science, Research and Arts Baden-Württemberg and the German Research Foundation (DFG) under Germany’s Excellence Strategy (CIBSS – EXC-2189 – Project ID 390939984). We gratefully acknowledge language editing by Anne Katrin Prowse. Funding by the Wissenschaftliche Gesellschaft Freiburg is gratefully acknowledged.

## Author declaration

All authors declare no conflict of interest.

## Data availability

Protein data have been deposited on ProteomeXchange.

## Supplementary Information

**Supplementary Figure S1.**
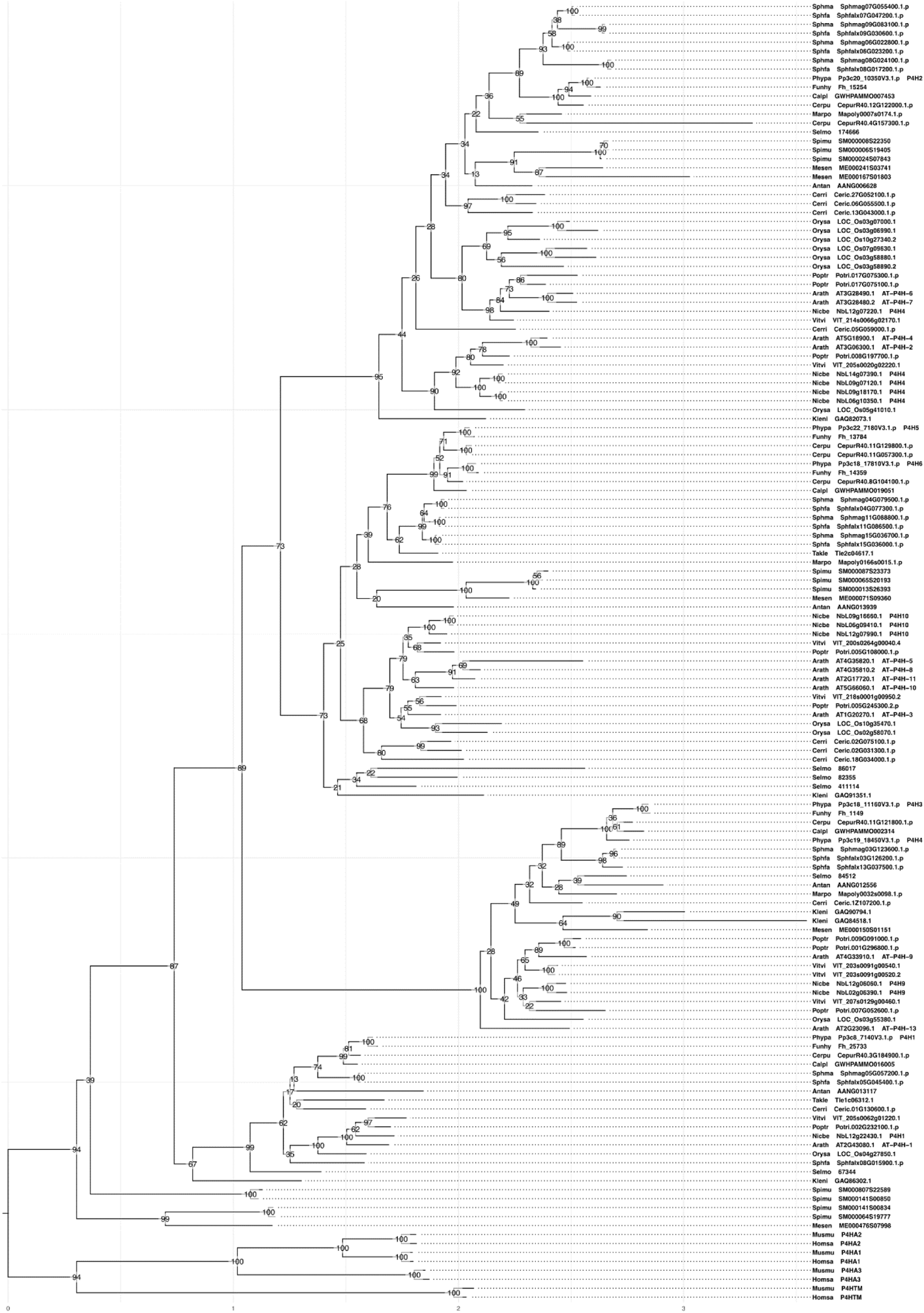
Phylogenetic reconstruction of the plant P4H family. Maximum-likelihood tree of a codon-aware multiple sequence alignment of P4H coding sequences annotated with all bootstrap support values, outgroup-rooted with mammal sequences

**Supplementary Figure S2.**
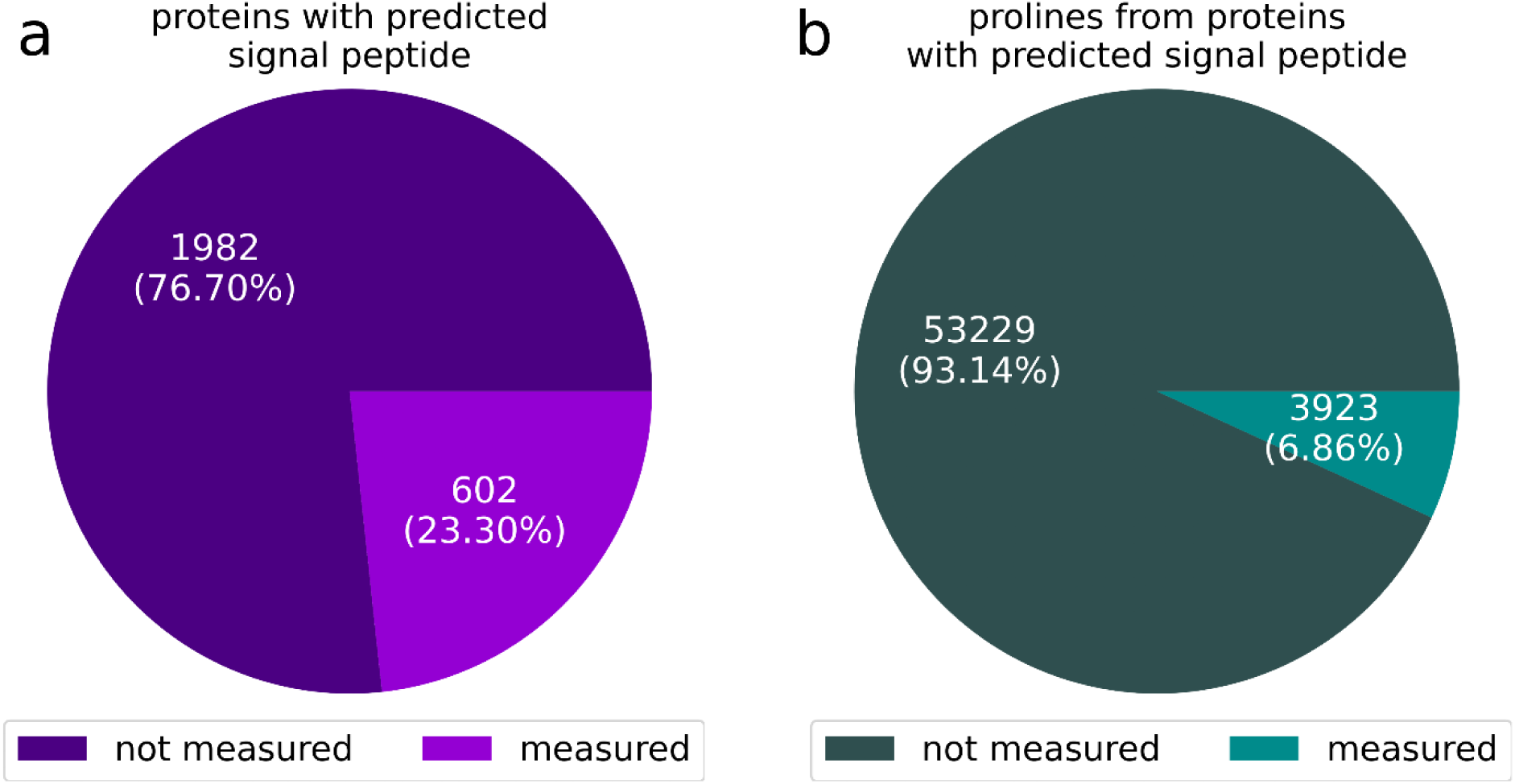
Overview of protein and proline coverage by the MS data. Depicted is the proportion of all secretory Physcomitrella proteins (proteins encoded by organelles are excluded) with a predicted signal peptide that were measured in any of the MS datasets (a) as well as the proportion of prolines from these proteins that were covered by a peptide in the MS data (b).

**Supplementary Figure S3.**
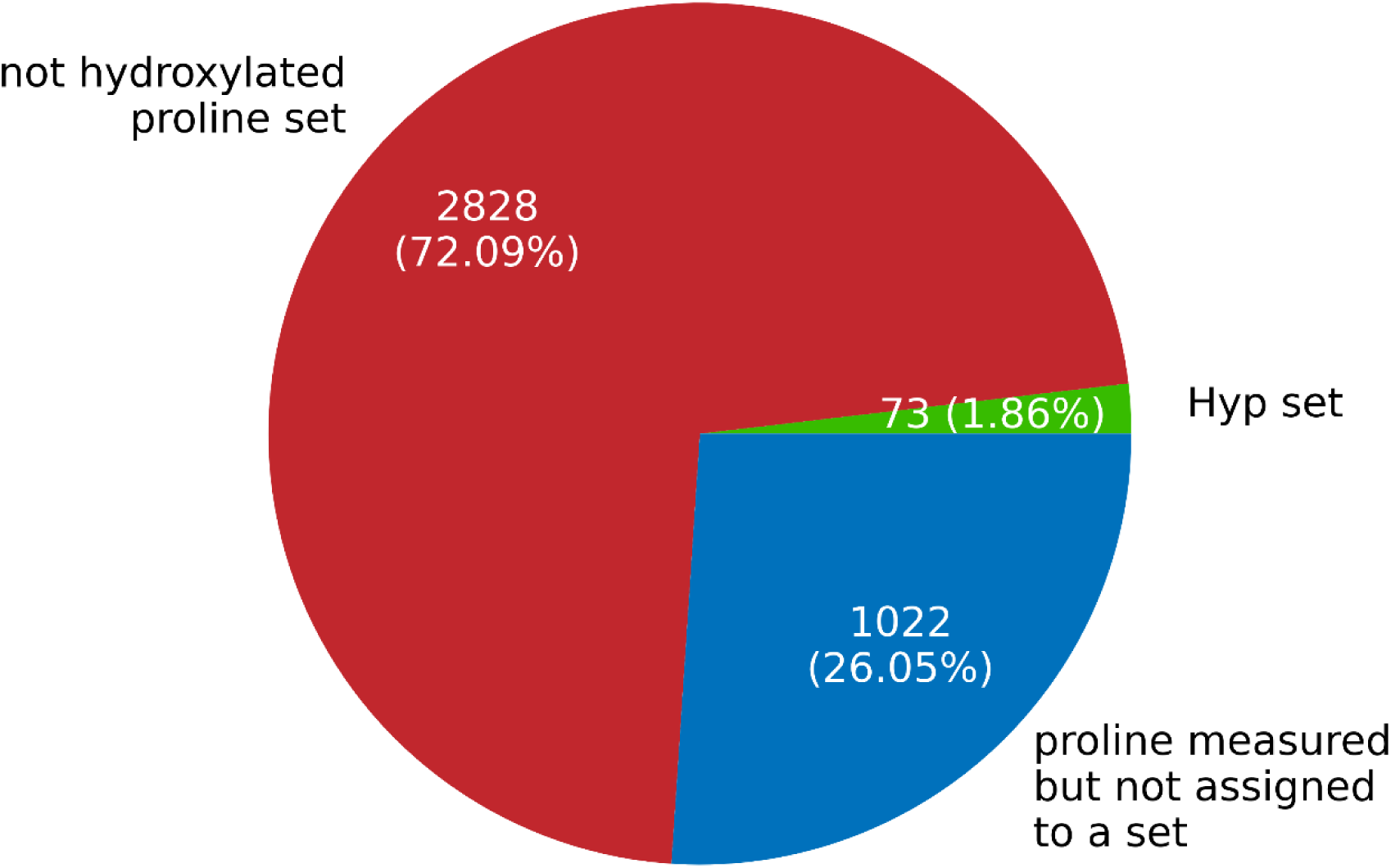
Overview of the hydroxylation status of the 3923 measured proline sites from secretory proteins with predicted signal peptide. Given are the percentage of prolines that were validated as Hyp (PTMProphet probability >0.7; green), the set of selected prolines that were not measured to be hydroxylated (PTMProphet probability for hydroxylation < 0.01; red) and the remaining prolines that were measured but not included e.g. because their hydroxylation status could not be unambiguously determined from the MS spectra (PTMProphet probability for hydroxylation > 0.01 and < 0.7; blue).

**Supplementary Table T1.**
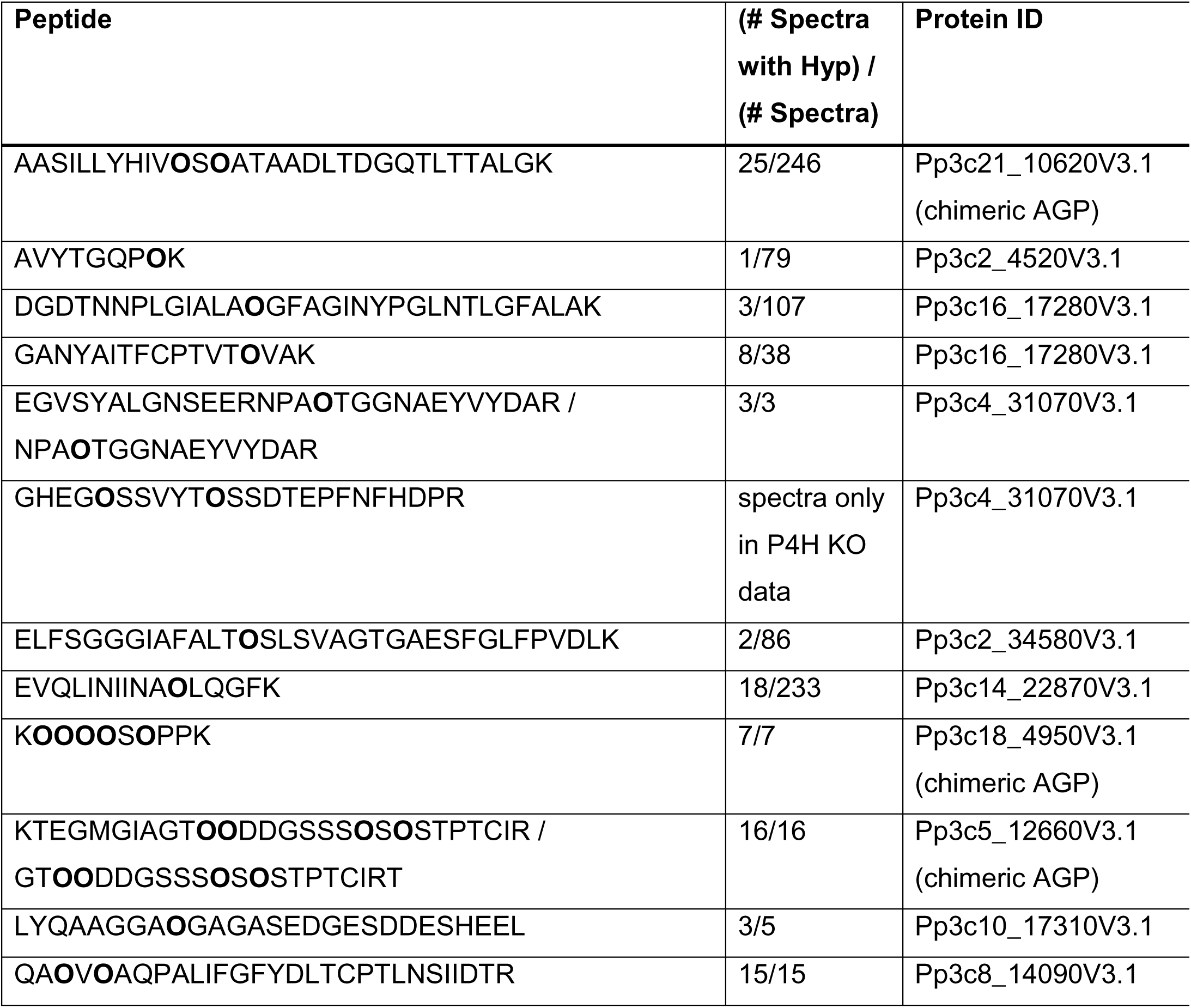
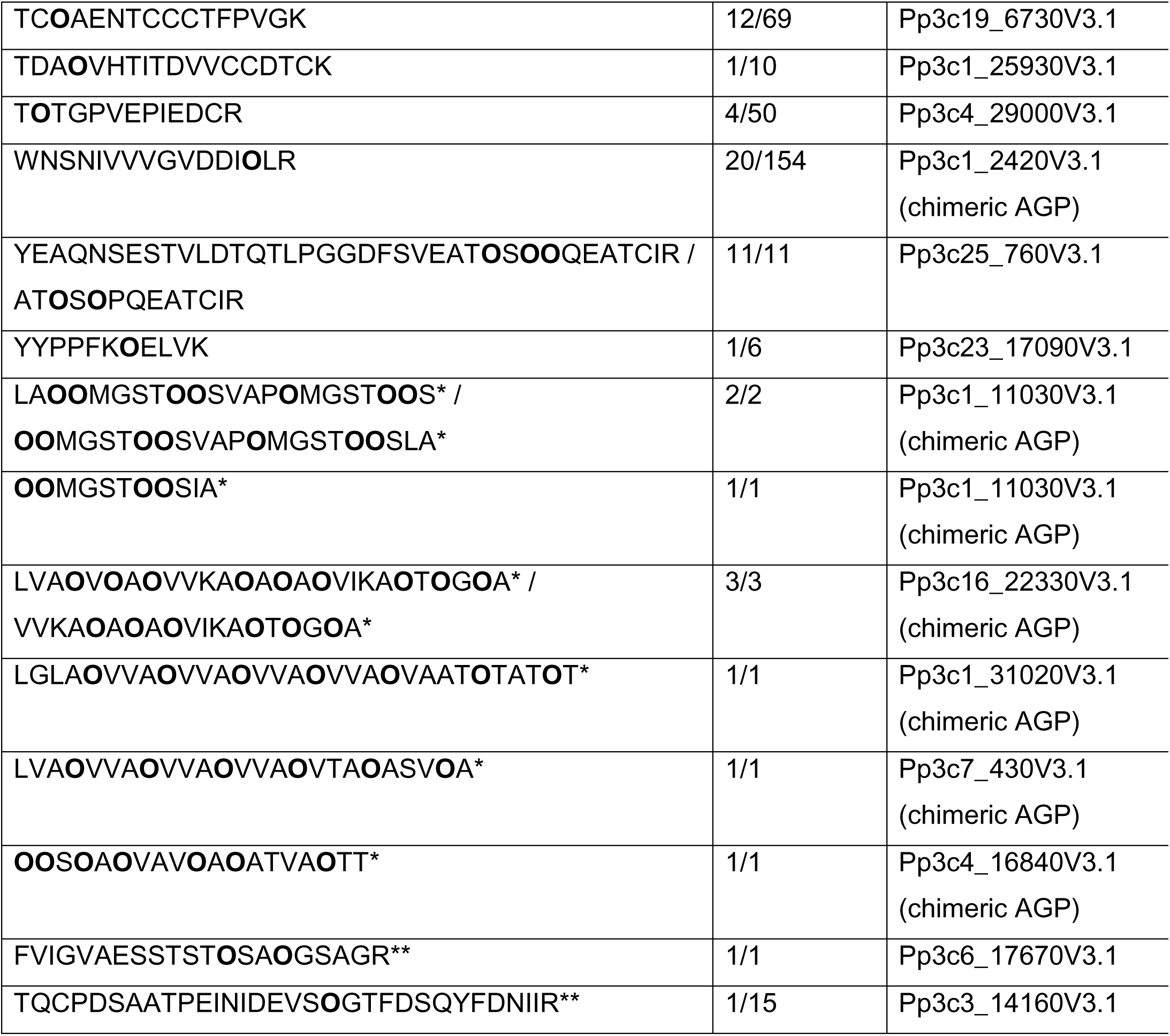
Overview of 26 Hyp-containing peptides collected from different mass spectrometry datasets. All possible prolines that were validated to be prolyl-hydroxylated in spectra of this peptide are written as an **O** in the AA sequence. In the second column the number of MS2 spectra of the peptide with any prolyl-hydroxylation and the number of total MS2 spectra of this peptide are given counted over all MS datasets used in this study excluding spectra from P4H KO lines. Spectra of the peptide GHEG**O**SSVYT**O**SSDTEPFNFHDPR were only identified in data of P4H KO lines (in P4H1 KO, P4H2 KO, P4H3 KO, P4H4 KO and P4H6 KO). Peptides whose MS2 spectra were exclusively measured in deglycosylated datasets are marked (*=chemically deglycosylated; **=galactosyltransferase double KO).

**Supplementary Figure S4.**
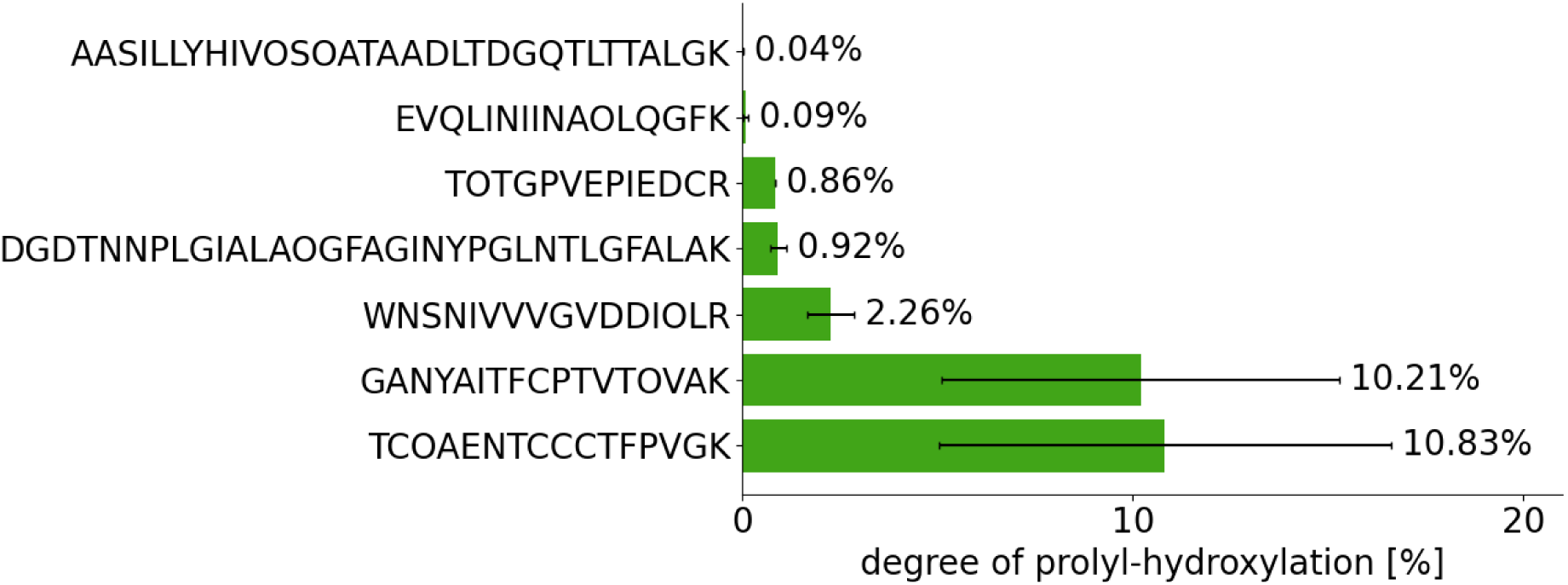
Estimation of the degree of hydroxylation of seven peptides from a quantified MS dataset. The mean percentage of the intensity of the summed prolyl-hydroxylated versions of a peptide with respect to the total intensity of all versions of the peptide was computed over three technical replicates each from two datasets.

**Supplementary Figure S5.**
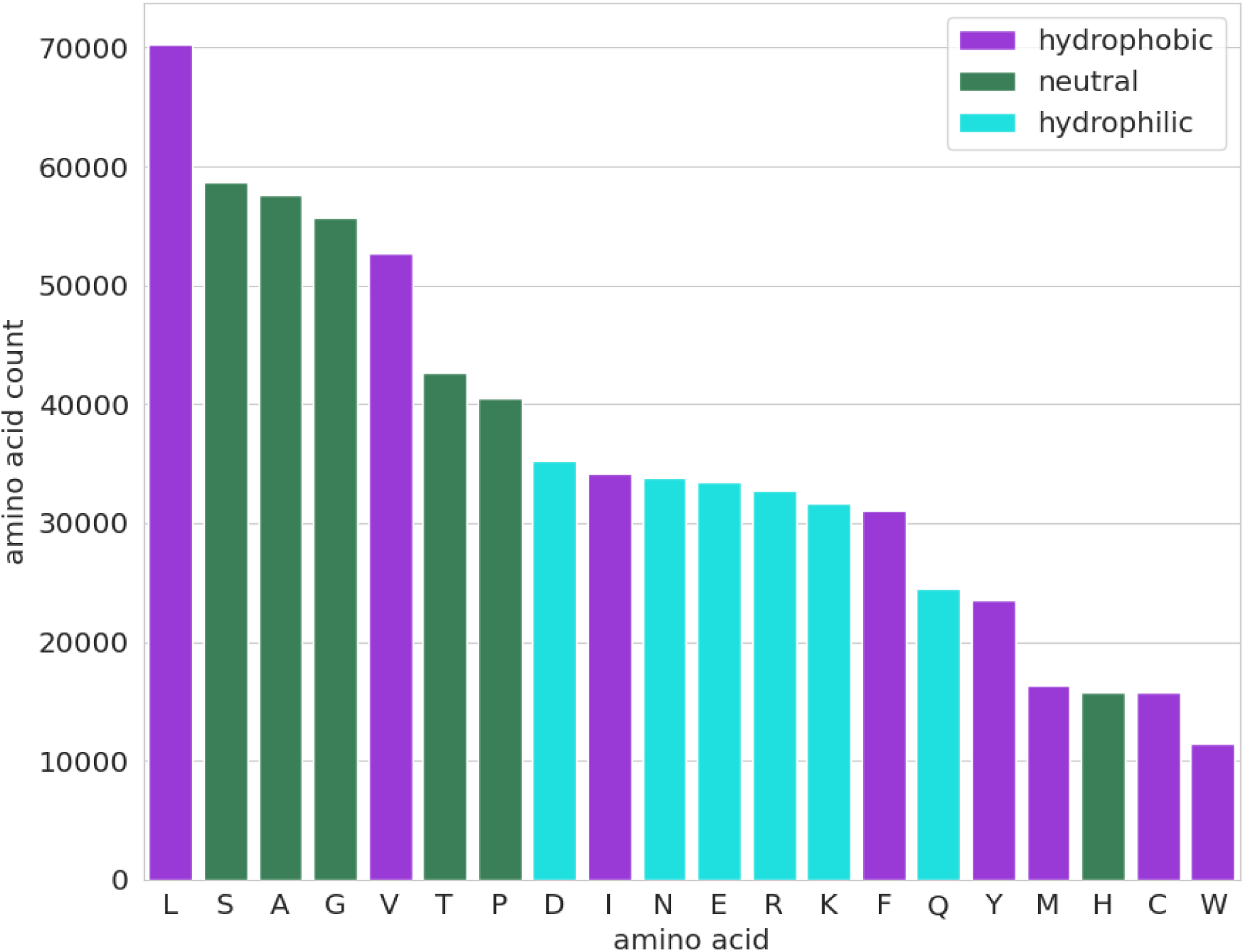
Overall amino acid distribution in the major isoforms of secretory Physcomitrella proteins with predicted signal peptide.

**Supplementary Table T2.**
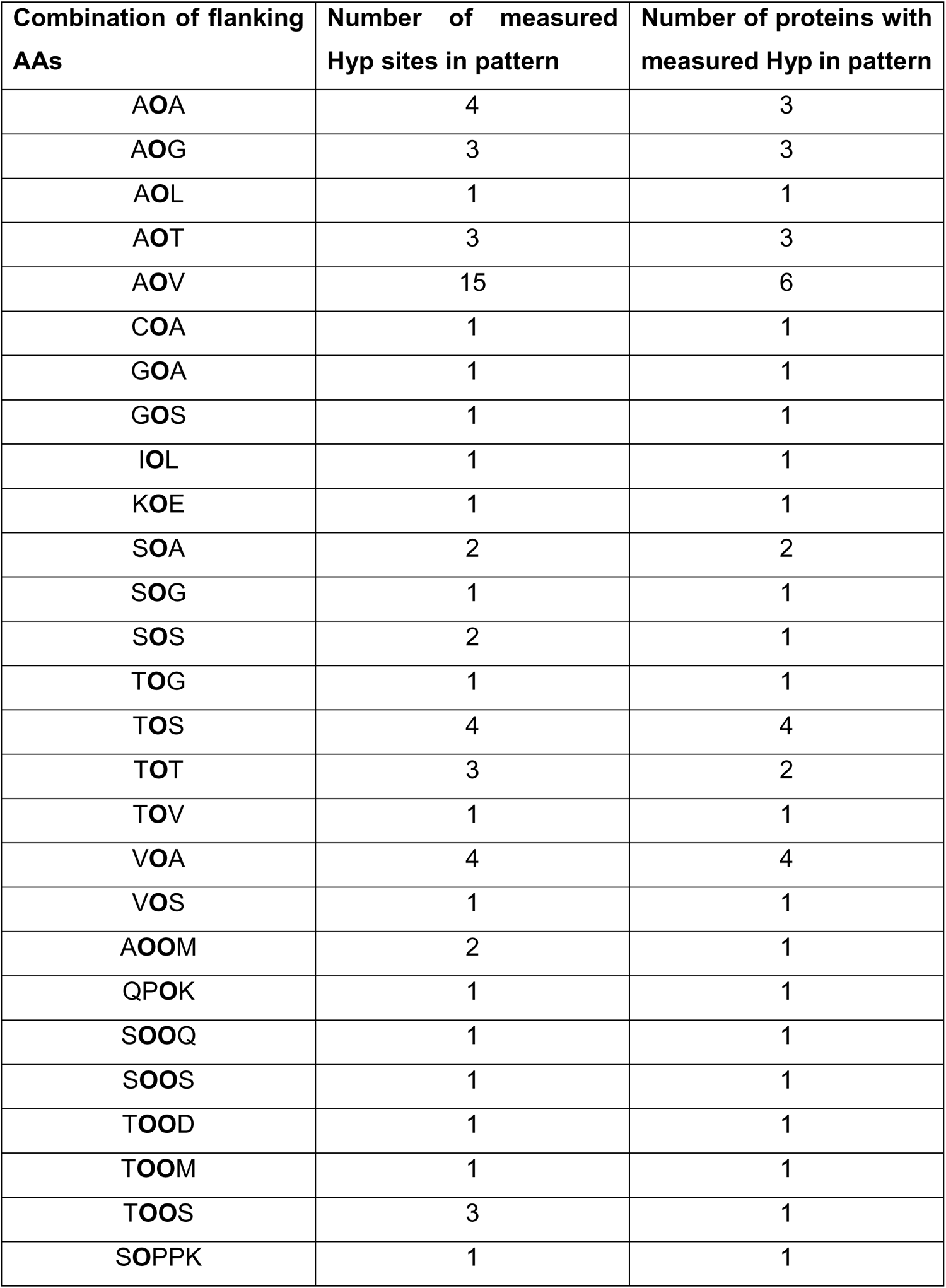
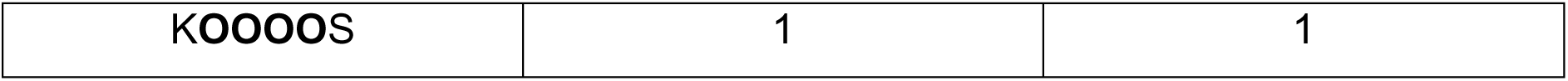
Combination of flanking amino acids around measured single Hyps or Hyp-containing proline blocks. Individual prolines that were at least once measured to be hydroxylated are noted as “O”.

**Supplementary Figure S6.**
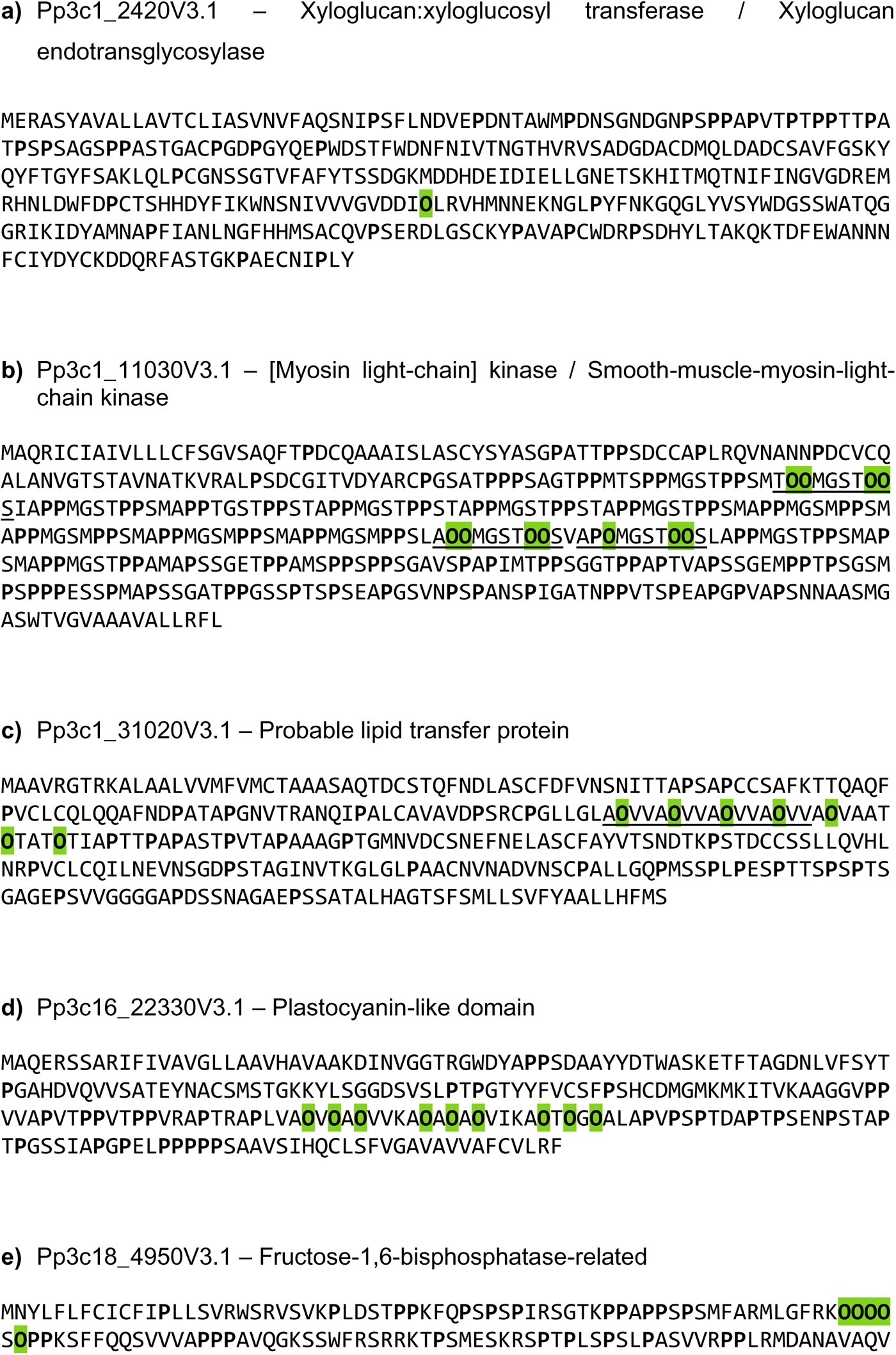
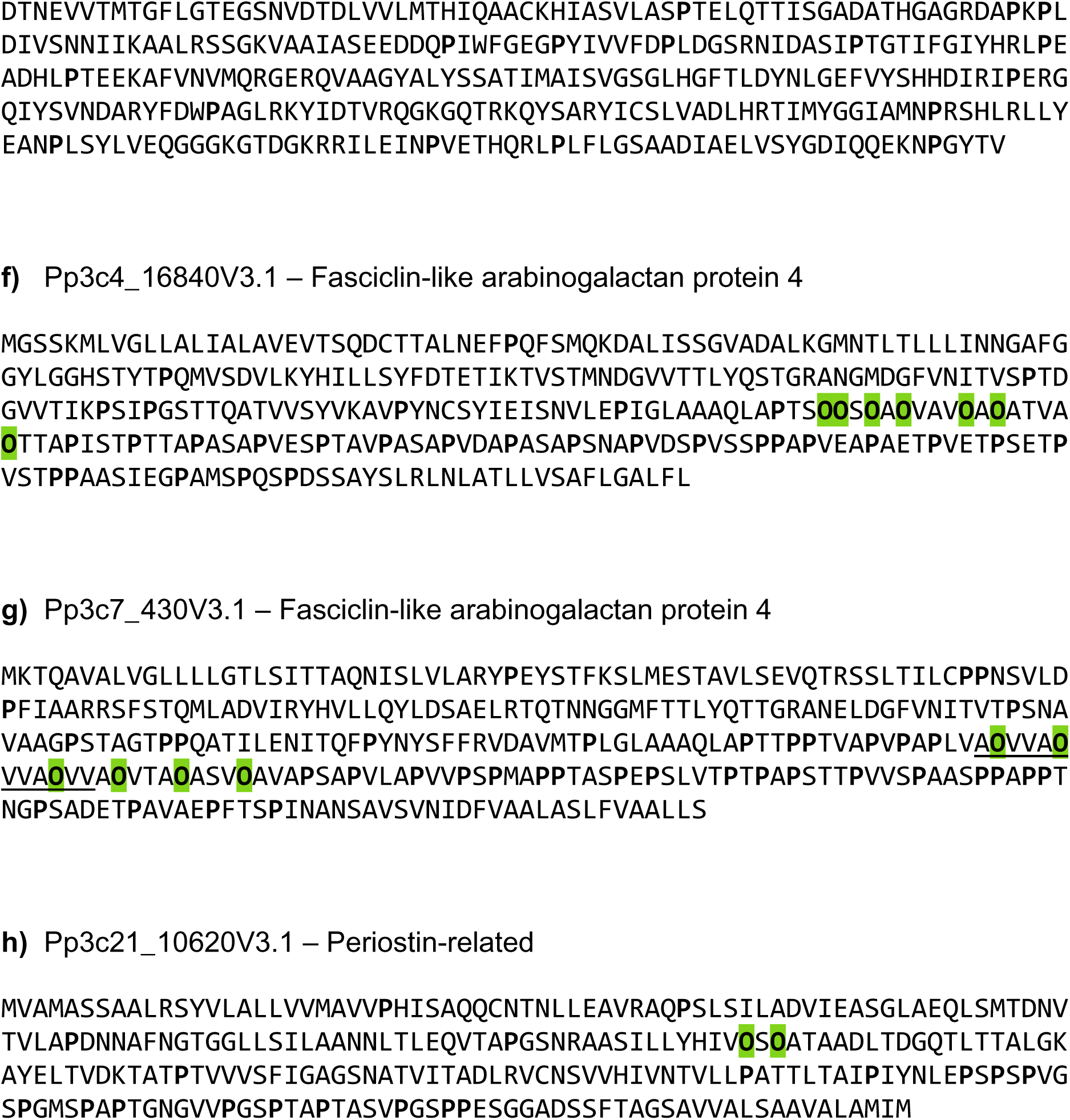
Amino acid sequences of chimeric AGPs with measured Hyps. All prolines are written in bold and measured Hyps (O) are highlighted in green. The repetitive Hyp-containing motifs A**O**VV and [A,T]**OO**MGST**OO** are underlined in the corresponding protein sequences.

**Supplementary Figure S7.**
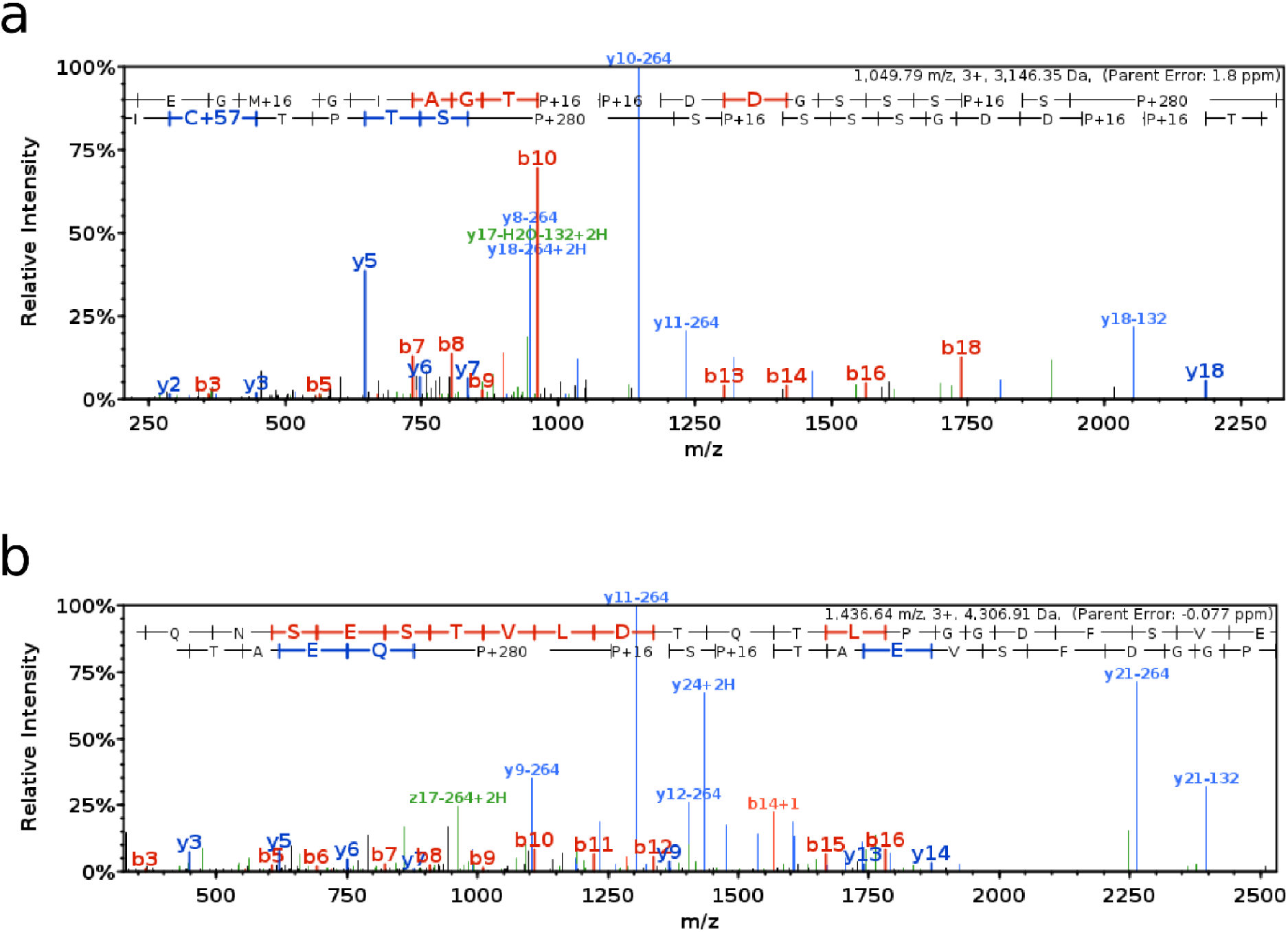
Fragment mass spectra from two peptides *O*-glycosylated with arabinose chains from pectinesterases. The peptides are (a) TEGMGIAGT**OO**DDGSSS**O**S**O**STPTCIR (Pp3c5_12660V3.1) and (b) YEAQNSESTVLDTQTLPGGDFSVEAT**O**S**OO**QEATCIR (Pp3c5_12660V3.1). y-ions with the neutral loss of one or two arabinose residues (-132 or -264) are present in both spectra. The image was generated in Scaffold 5 (Proteome Software, Inc., Portland, OR 97219, USA).

**Supplementary Figure S8.**
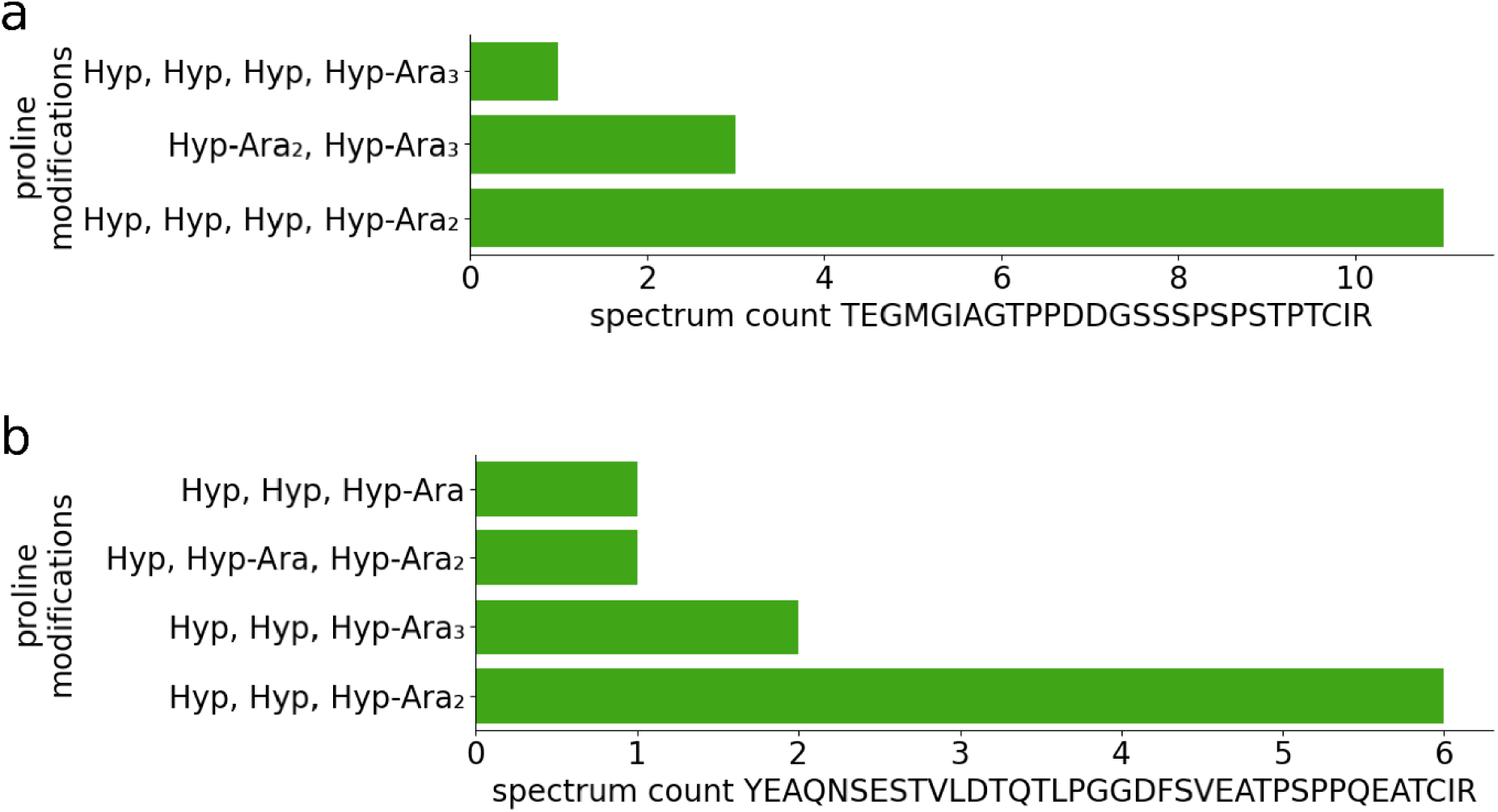
Modified versions of peptides from two secretory pectinesterases that were *O*-glycosylated with arabinose residues. Depicted are the number of spectra of the unmodified, prolyl-hydroxylated or the arabinosylated versions of the peptides TEGMGIAGT<u>PP</u>DDGSSS<u>P</u>S<u>P</u>STPTCIR (Pp3c5_12660V3.1) (a) and YEAQNSESTVLDTQTLPGGDFSVEAT<u>P</u>S<u>PP</u>QEATCIR (Pp3c5_12660V3.1) (b). Prolines modified with prolyl-hydroxylation (Hyp: +15.99) or additional *O*-glycosylation with arabinose residues (Hyp-Ara: +148.04; Hyp-Ara₂: +280.08; Hyp-Ara₃: +412.12) are underlined. Peptides with missed cleavage sites were considered. The spectra are counted over several MS measurements.

**Supplementary Figure S9.**
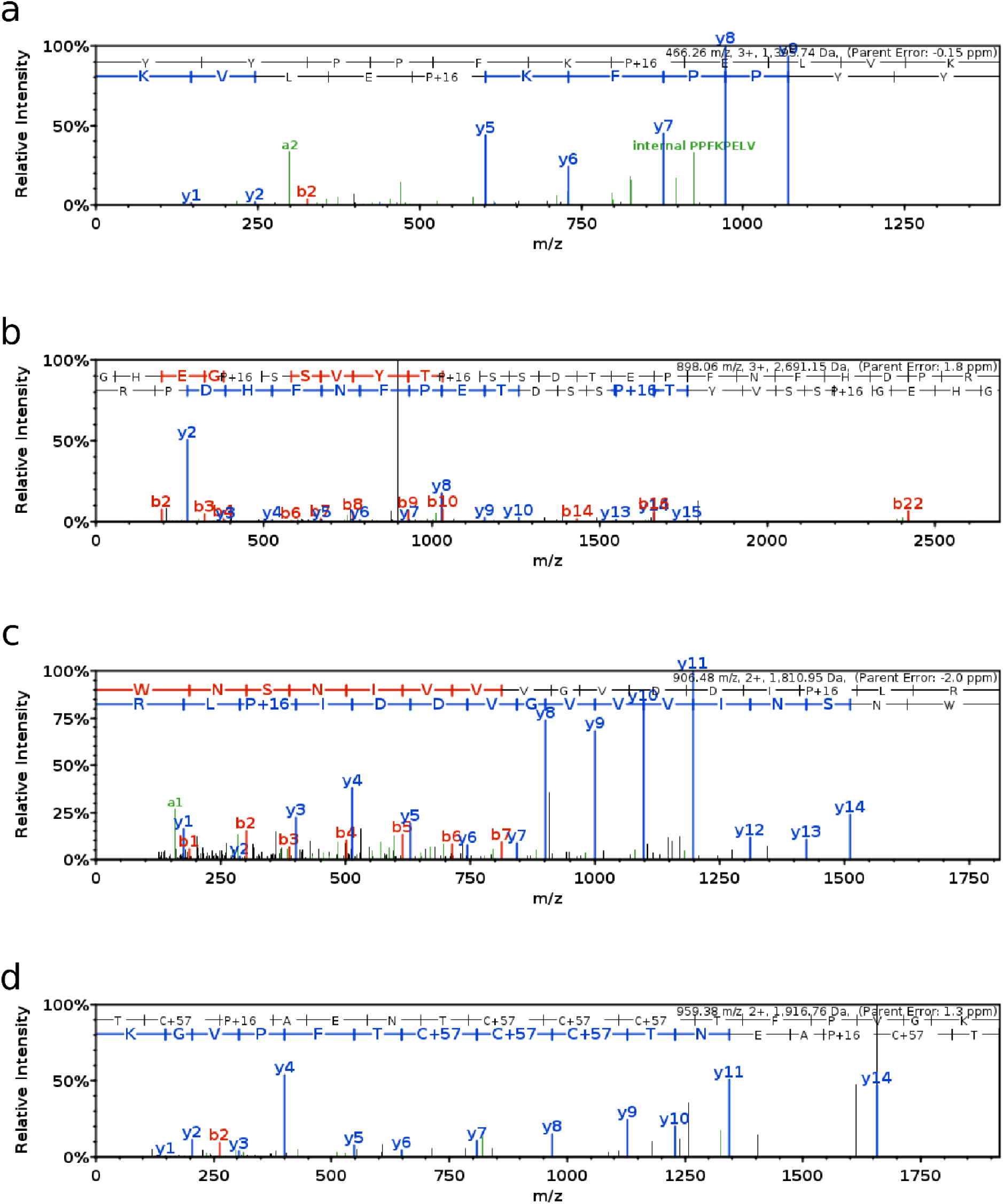
Fragment mass spectra from four Hyp-containing peptides that were not predicted to be prolyl-hydroxylated by three methods. The peptides are: (a) YYPPFK**O**ELVK, (b) GHEG**<u>O</u>**SSVYT**O**SSDTEPFNFHDPR (underlined Hyp was not predicted), (c) WNSNIVVVGVDDI**O**LR and (d) TC**O**AENTCCCTFPVGK. The images were generated in Scaffold 5 (Proteome Software, Inc., Portland, OR 97219, USA).

**Supplementary Figure S10.**
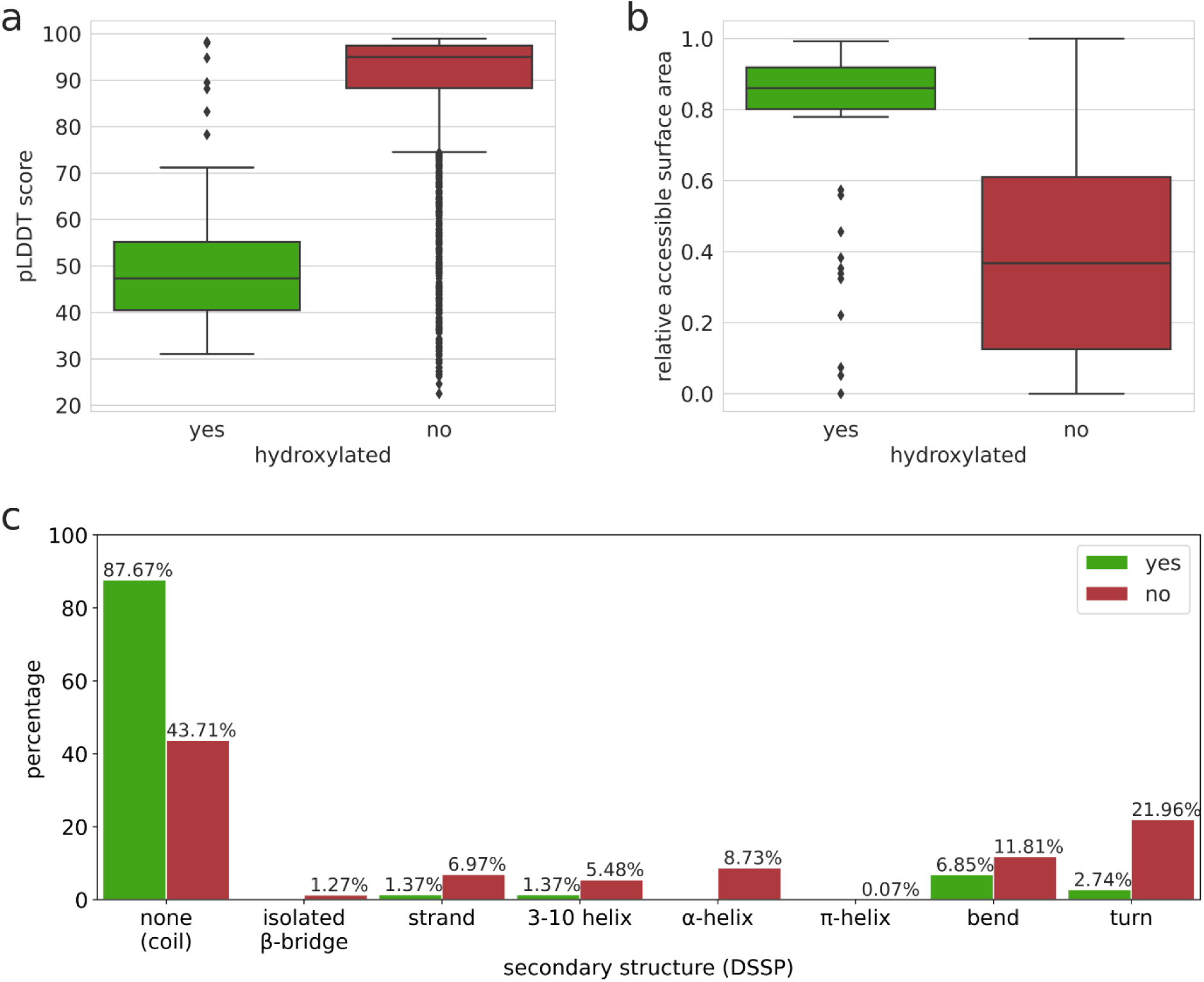
Structural environment of Hyp sites and not hydroxylated proline sites in protein structure models. The models were downloaded from the AlphaFold Protein Structure Database (Varadi et al., 2022). Depicted is the distribution of the pLDDT confidence scores (0=minimum quality, 100=maximum quality) of Hyp residues and proline residues that were not measured to be hydroxylated (a), their relative accessible surface area (0=completely buried within protein structure, 1=fully exposed to solvent) (b) as well as secondary structure elements containing the respective residues (c). The relative accessible surface area and the secondary structure elements were assigned with DSSP.

**Supplementary Figure S11.**
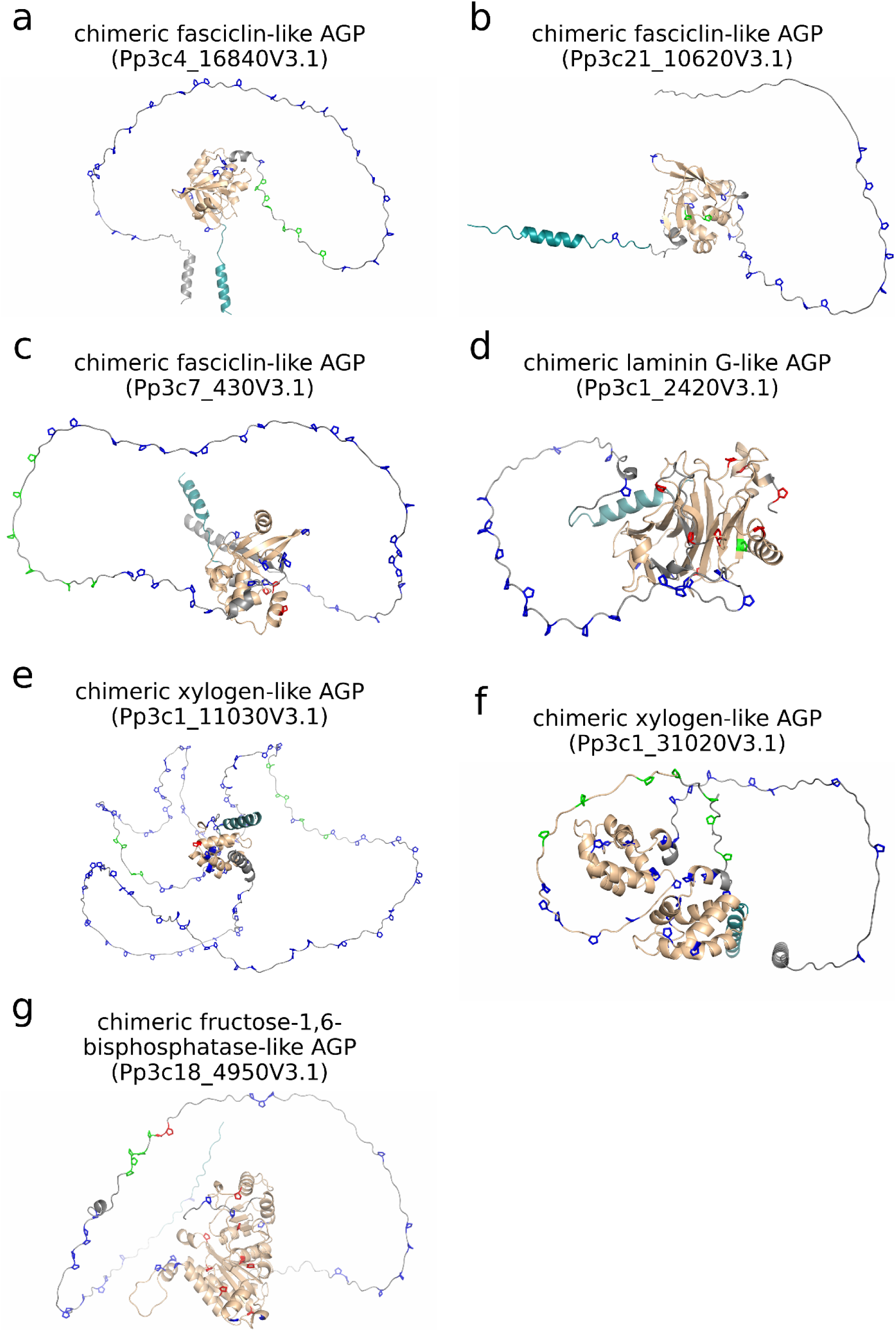
Three-dimensional structure models of HRGPs with validated Hyps. Validated Hyp sites (green) and proline sites not measured to be hydroxylated (red) are highlighted in the models downloaded from the AlphaFold Protein Structure Database (Varadi et al., 2022). All remaining prolines are coloured blue. For these, no certain information about their hydroxylation status could be gained from the MS data. The structures are from chimeric AGPs as predicted by Ma et al. (2017). The following Pfam domains as given by Phytozome (v13; Goodstein et al., 2012) are coloured in light brown: a), b) and c) fasciclin domain (PF02469); d) glycosyl hydrolases family 16 (PF00722) and Xyloglucan endo-transglycosylase (XET) C-terminus (PF06955); e) and f) probable lipid transfer (PF14368); g) fructose-1-6-bisphosphatase, N-terminal domain (PF00316). Signal peptides (SignalP 5, Almagro Armenteros et al., 2019a) are coloured in turquoise.

**Supplementary Figure S12.**
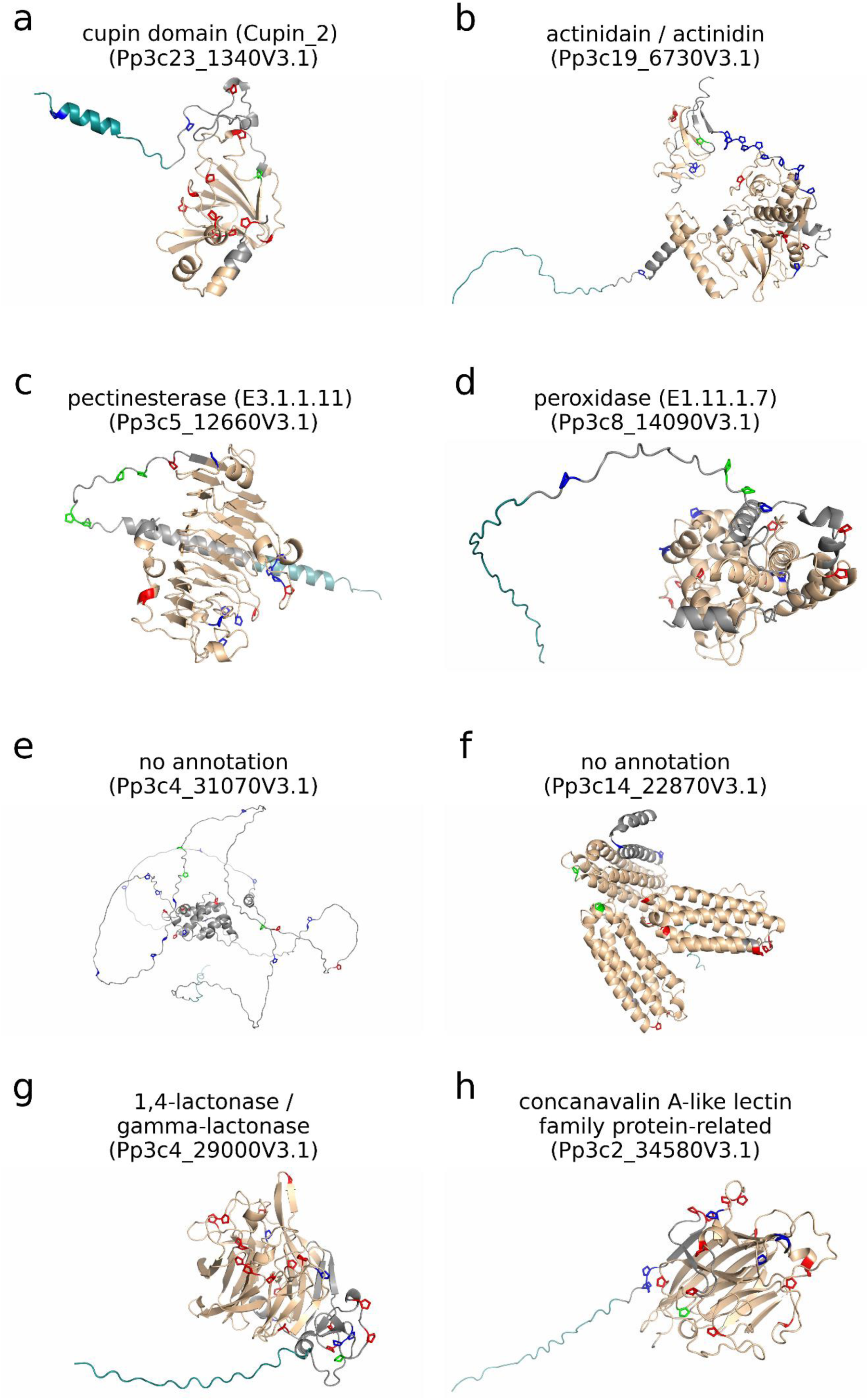
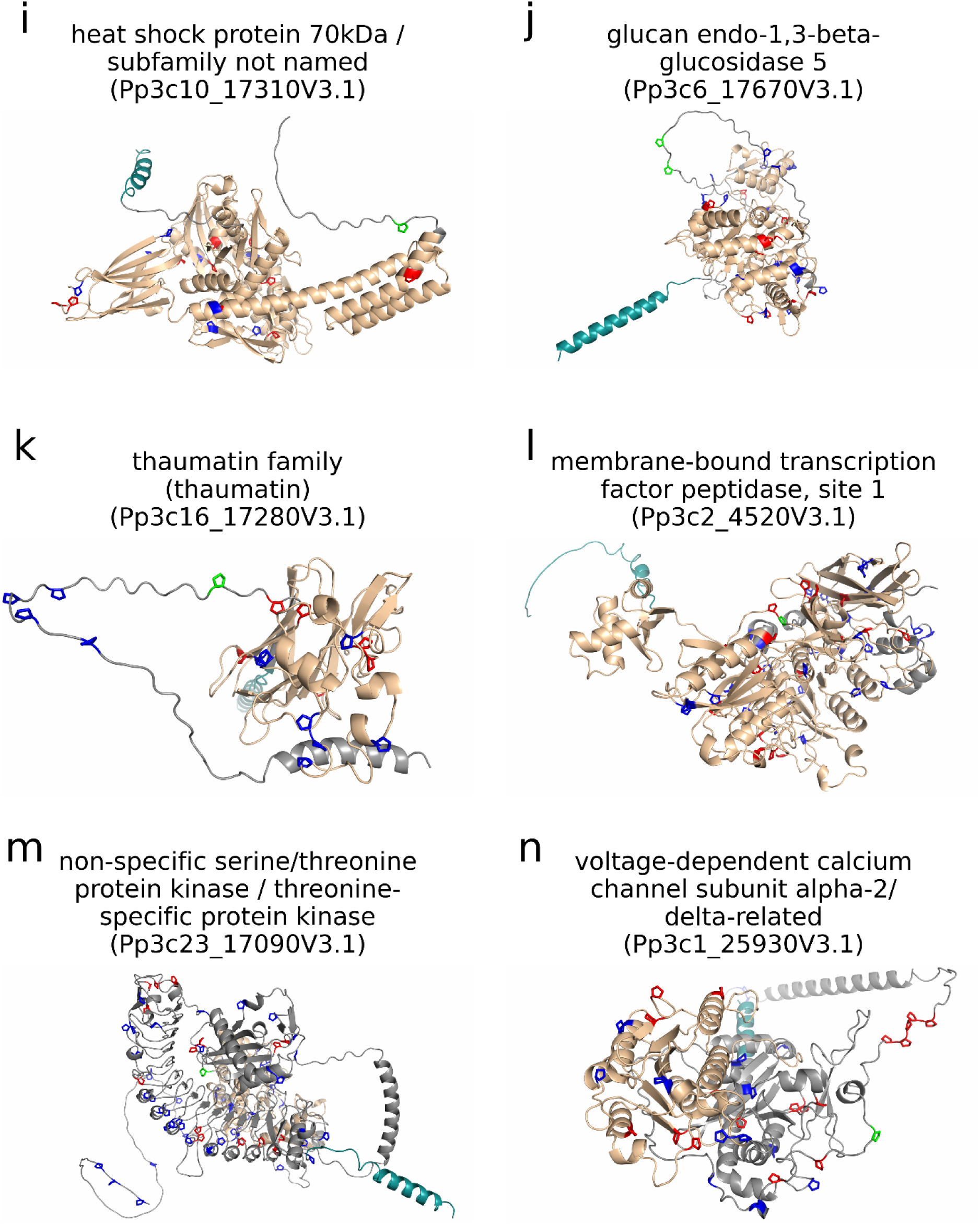
Three-dimensional structure models of proteins with validated Hyp sites that were not predicted to be HRGPs. Validated Hyp sites (green) and proline sites not measured to be hydroxylated (red) are highlighted in the models downloaded from the AlphaFold Protein Structure Database (Varadi et al., 2022). All remaining prolines are coloured blue. For these, no certain information about their hydroxylation status could be gained from the MS data. The annotation is based on Phytozome (v13; Goodstein et al., 2012) and the following Pfam domains as given by Phytozome are coloured in light brown: a) cupin (PF00190); b) granulin (PF00396), papain family cysteine protease (PF00112) and cathepsin propeptide inhibitor domain (I29; PF08246); c) pectinesterase (PF01095); d) peroxidase (PF00141); f) family of unknown function (PF17615); g) SMP-30/gluconolactonase/LRE-like region (PF08450); h) legume lectin domain (PF00139); i) Hsp70 protein (PF00012); j) glycosyl hydrolases family 17 (PF00332) and X8 domain (PF07983); k) thaumatin family Pfam entry (PF00314); l) subtilase family (PF00082), peptidase inhibitor I9 (PF05922) and fibronectin type-III domain (PF17766); m) protein kinase domain (PF00069); n) von Willebrand factor type A domain (PF13519). Signal peptides (SignalP 5, Almagro Armenteros et al., 2019a) are coloured in turquoise.

**Supplementary Figure S13.**
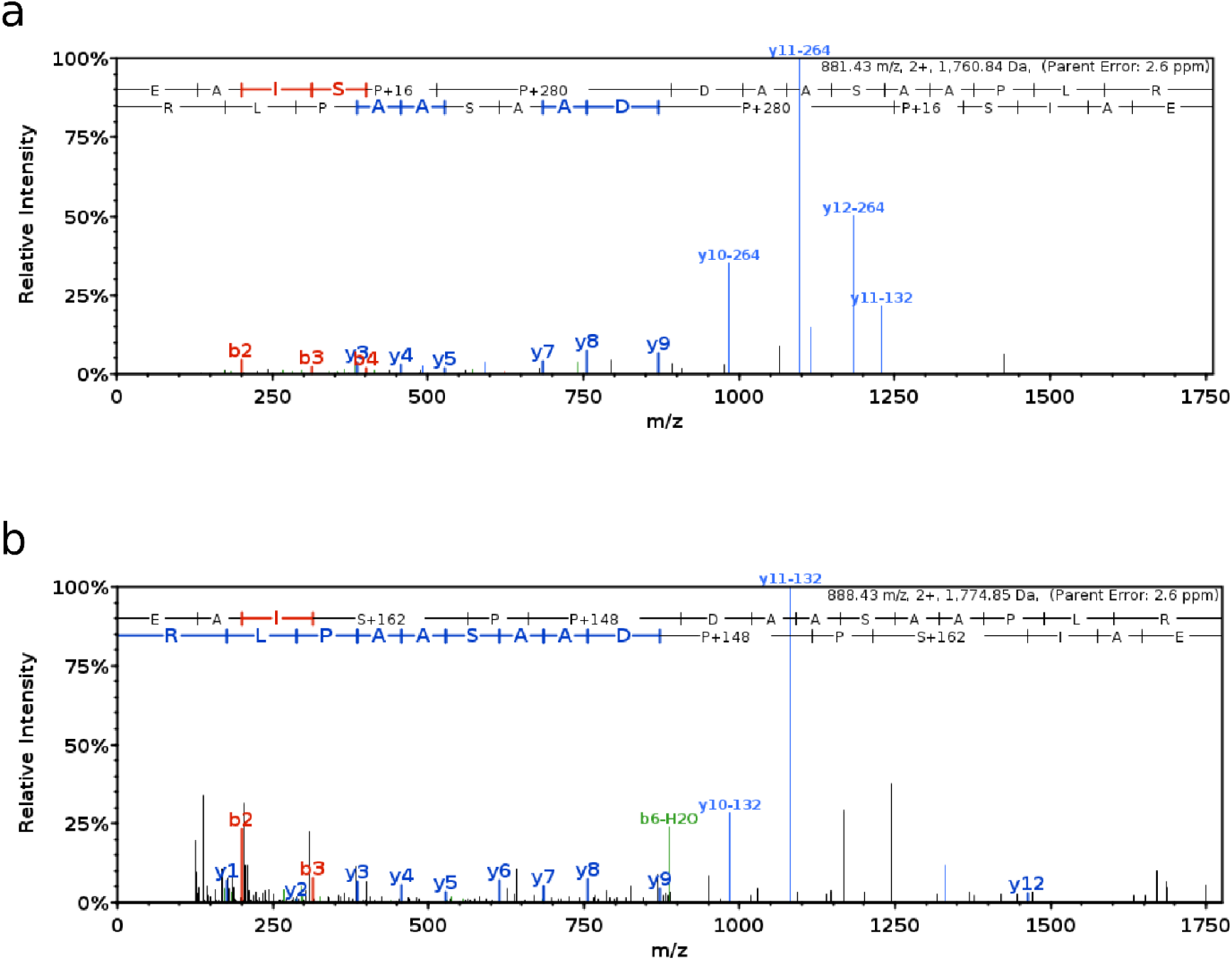
Fragment mass spectra from the EPO peptide EAISOODAASAAPLR with hydroxylated prolines and *O*-glycosylation. An example for *O*-glycosylation with two arabinose residues (a) and *O*-glycosylation of the first serine with a hexose are given (b). In (a) the y-ions y10, y11 and y12 are present with the neutral loss of one or two arabinose residues (-132 or -264). In (b) the y-ions y10, y11 and y12 are present with the neutral loss of one arabinose (-132). The image was generated in Scaffold 5 (Proteome Software, Inc., Portland, OR 97219, USA).

**Supplementary Figure S14.**
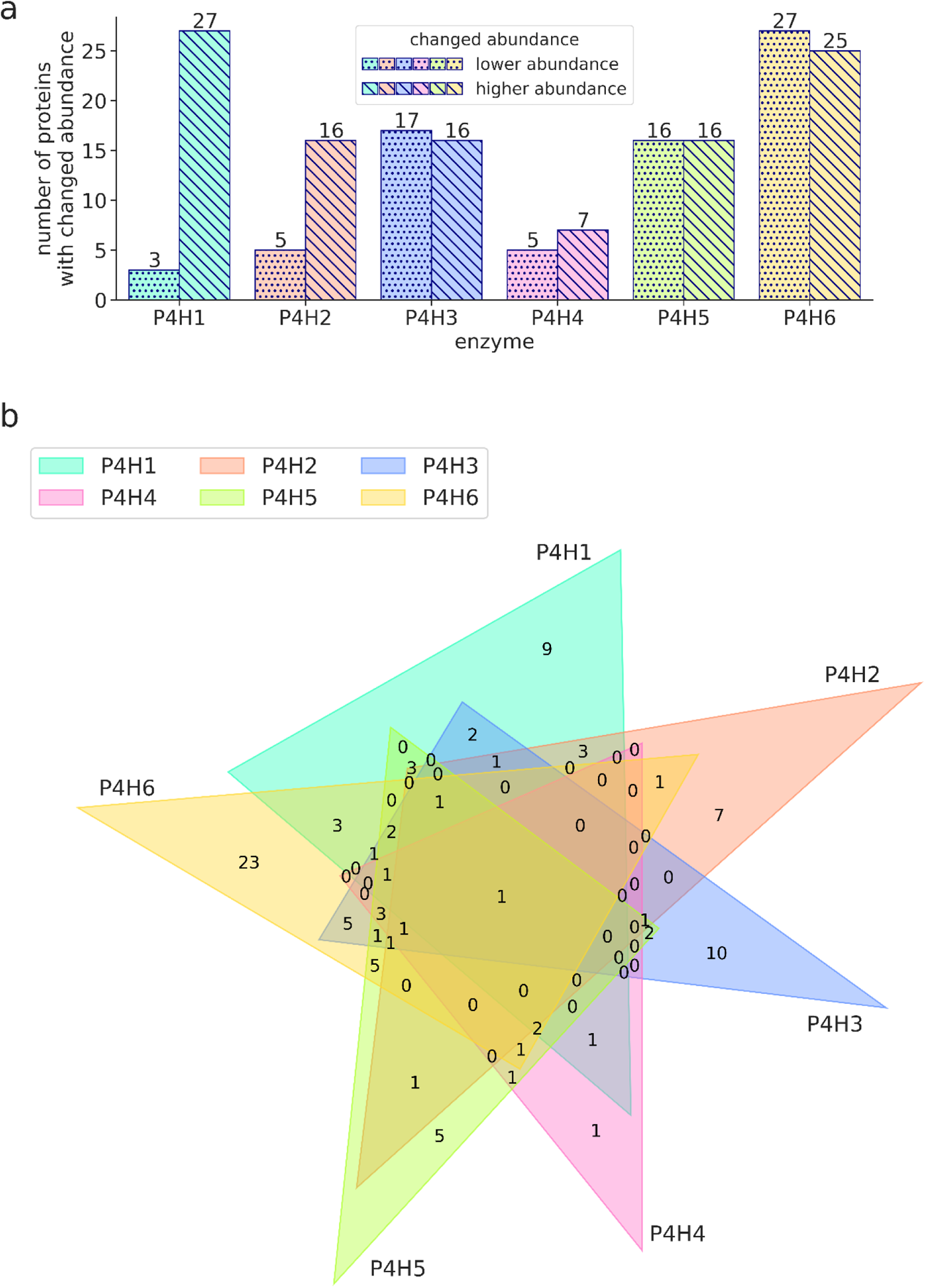
Proteins with changed abundance in P4H1-P4H6 single KO lines compared to the maternal line. The proteins are from MS measurements of samples enriched with secretory proteins using two different protocols. Significant changes in the intensity light/heavy ratio (intensity in P4H KO line/intensity in maternal line) of proteins were determined using a *t*-test (*P* adjusted < 0.05; |log2 light/heavy ratio|>1). Only proteins with predicted signal peptide are depicted and results from both protocols were combined. (a) Number of proteins with increased or decreased abundance in the P4H KO lines compared to the maternal line. (b) Overlap of proteins with changed abundance from the P4H KO lines compared to the parental line.

**Supplementary Table T3.** HRGPs with changed abundance in single P4H KO lines compared to maternal line. MS samples enriched with secretory proteins were obtained using two different protocols. Given are the average log2 light/heavy intensity ratios (intensity in P4H KO line/intensity in maternal line) of the protein in the P4H KO lines compared to the maternal line from three technical replicates. Values are only included in the table if the protein was present in all three replicates and the change was significant (*t*-test with *P* < 0.05; |log2 light/heavy ratio| > 1). Otherwise the cell is left empty. In the P4H4 KO line no HRGP had a significantly changed abundance.

| **Protein ID and annotation** | **HRGP class** | **P4H1 KO** | **P4H2 KO** | **P4H3 KO** | **P4H5 KO** | **P4H6 KO** |
| --- | --- | --- | --- | --- | --- | --- |
| Xyloglucan endotransglycosylase (Pp3c1_2420V3.1) | Chimeric AGP | 2.50 |  |  |  | -2.57 |
| Fasciclin-like arabinogalactan protein 4 (Pp3c26_5590V3.1) | Chimeric AGP | 5.39 | 6.38 | 5.46 | 5.94 | 7.62 |
| Fasciclin-like arabinogalactan protein 4 (Pp3c4_16840V3.1) | Chimeric AGP | 12.54 | 9.08 | 5.43 |  |  |
| Plastocyanin-like domain (Pp3c16_22330V3.1) | Chimeric AGP |  |  |  |  | 5.16 |

**Supplementary Table T4.**
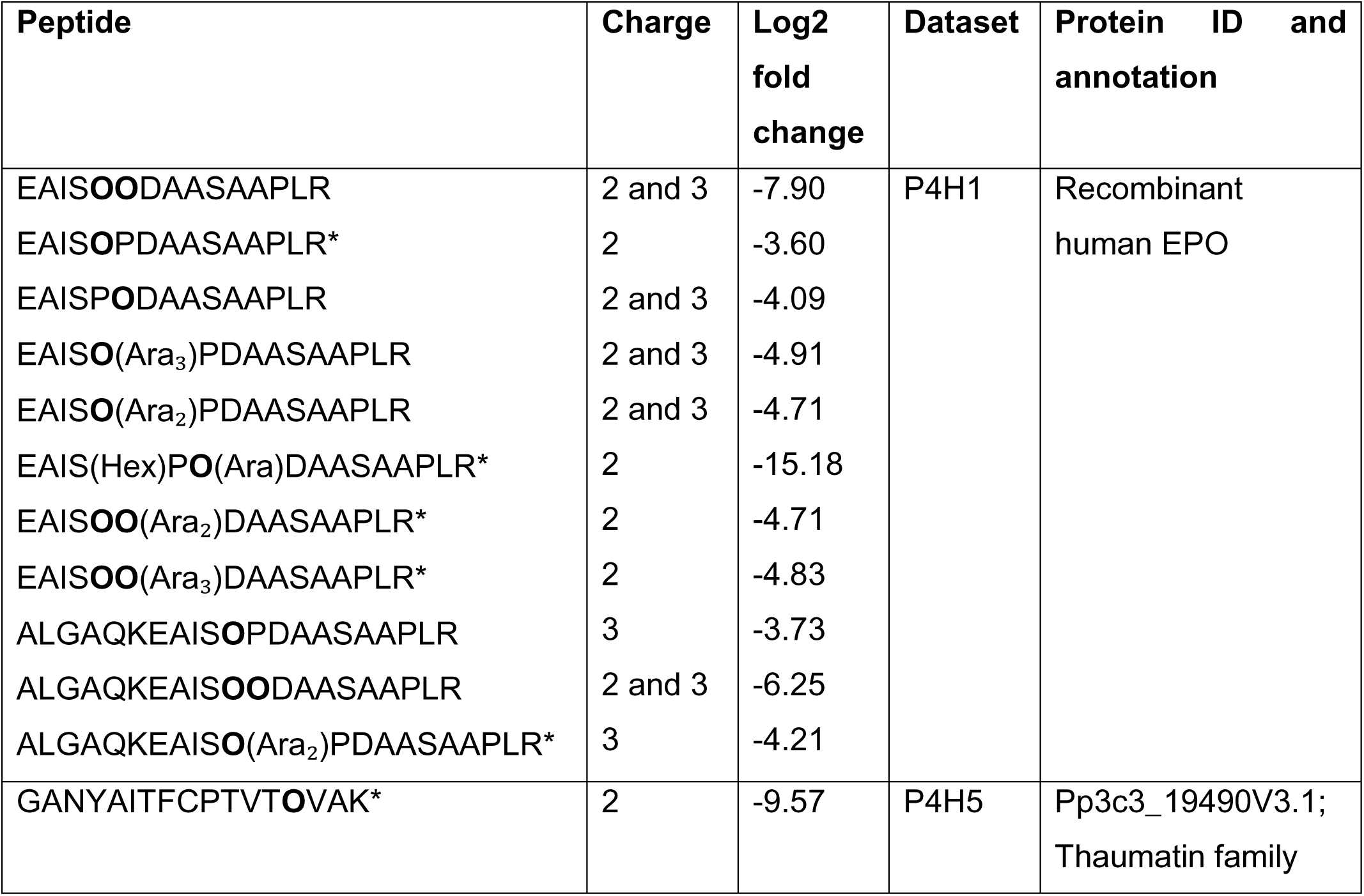
Prolyl-hydroxylated peptides with reduced abundance in P4Hs single KO lines compared to maternal line. The peptides are from MS measurements using three replicates from two protocols to enrich secretory proteins. Light/heavy intensity ratios (intensity in P4H KO line/intensity in maternal line) of peptides from the same protein were normalized by the median light/heavy ratio of all peptides from the respective protein. Significant changes of the light/heavy ratio of peptides were determined with a *t*-test (*P* < 0.05 and |log2 fold change| > 1) and, if present, changes in the level of the unmodified peptide were also considered. Only Hyp-containing peptides with significantly reduced abundance in the P4H KO lines were selected. Peptides marked with * were only measured in two of the replicates. If the abundance of a peptide with the same modifications was significantly reduced in data from both protocols or if the reduction was significant for multiple charges, the average log2 fold change is given. Peptides with hexose (Hex: + 162.05), mono-arabinosylation (Ara: + 148.04), di-arabinosylation (Ara₂: + 280.08) or tri-arabinosylation (Ara₃: + 412.12) are present.

**Supplementary Figure S15.**
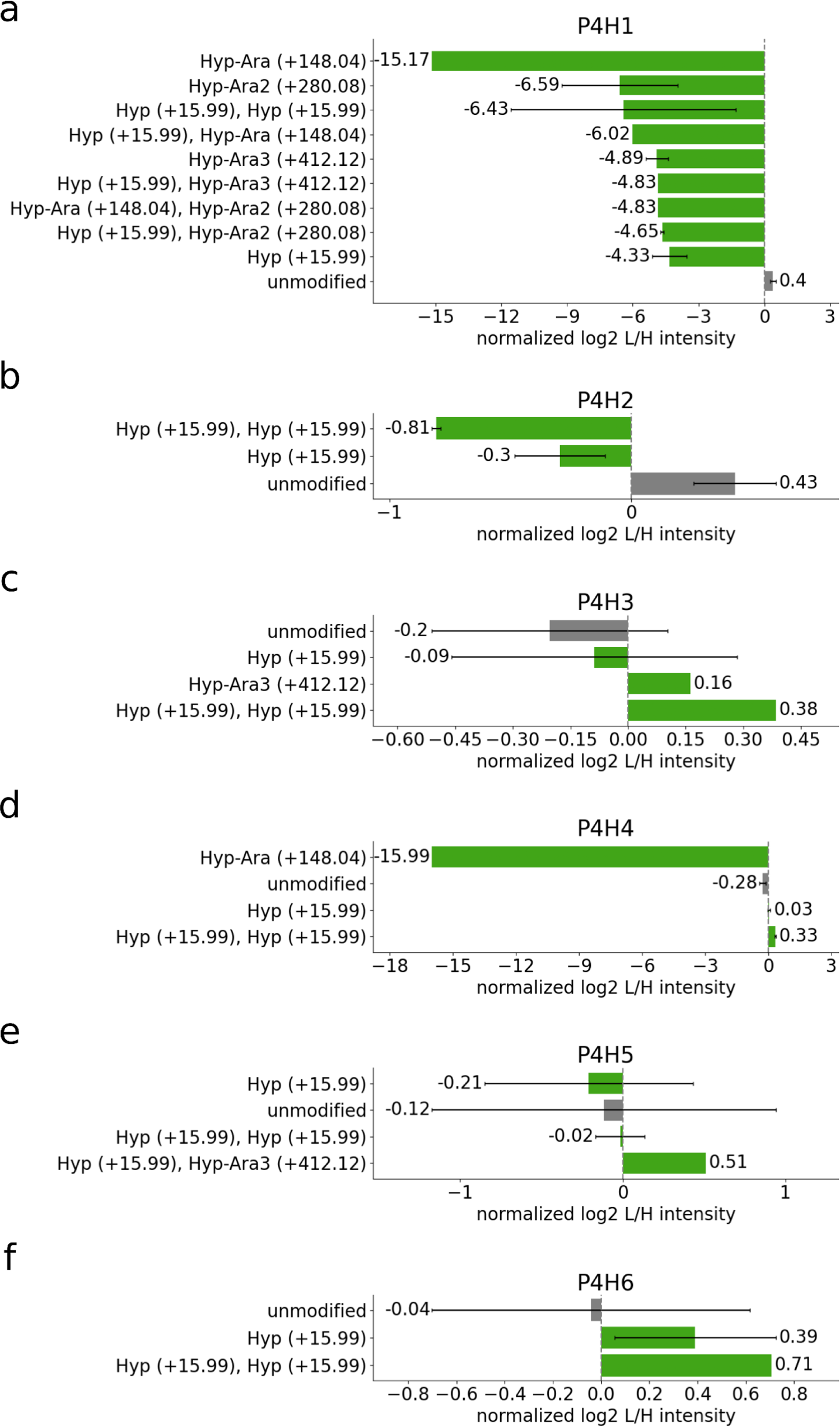
Abundance ratio of modified and unmodified versions of the EPO peptide EAISPPDAASAAPLR in P4H KO lines compared to the maternal line. The peptides are from MS measurements using three replicates from two protocols to enrich secretory proteins. Log2 light/heavy intensity ratios within a replicate were normalized by the median light/heavy intensity ratios of all EPO peptides. For each bar the mean of the log2 light/heavy intensity ratios (=intensity in P4H KO line/intensity in maternal line) of all peptides with the specified combination of proline modifications was taken. (a) P4H1 KO/maternal line, (b) P4H2 KO/maternal line, (c) P4H3 KO/maternal line, (d) P4H4 KO/maternal line, (e) P4H5 KO/maternal line, (f) P4H6 KO/maternal line. The strong reduction of Hyp-Ara in (d) is based on the L/H ratio of only one peptide that carried an additional hexose on the first serine.

**Supplementary Figure S16.**
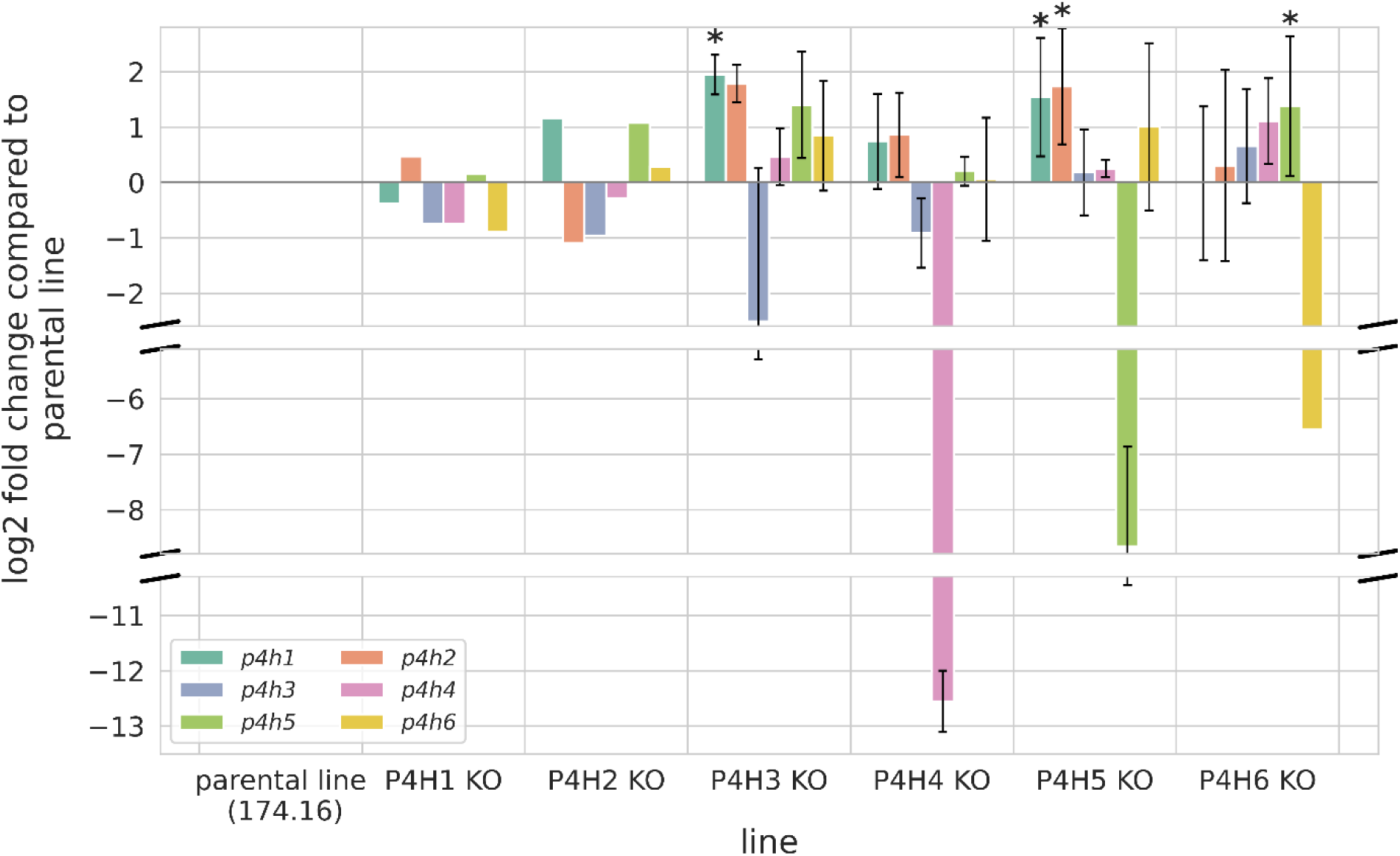
Average expression of the six different P4H genes in P4H KO lines. The expression is normalized against the maternal line and the log2 fold change is displayed. Following lines were tested using biological replicates: P4H1 KO and P4H2 KO: n=2; P4H3 KO, P4H4 KO, P4H5 KO, P4H6 KO and the maternal line (174.16): n=3. Following genes were tested: *p4h1*, *p4h2*, *p4h3*, *p4h4*, *p4h5*: n= 3, *p4h6*: n=1. The standard deviation of the average log2 fold expression is displayed. Significant fold changes (*P* < 0.05) are marked with (*). Statistical analysis was performed with GraphPad Prism 8 using an ANOVA with Durentt‘s test.

**Supplementary Figure S17.**
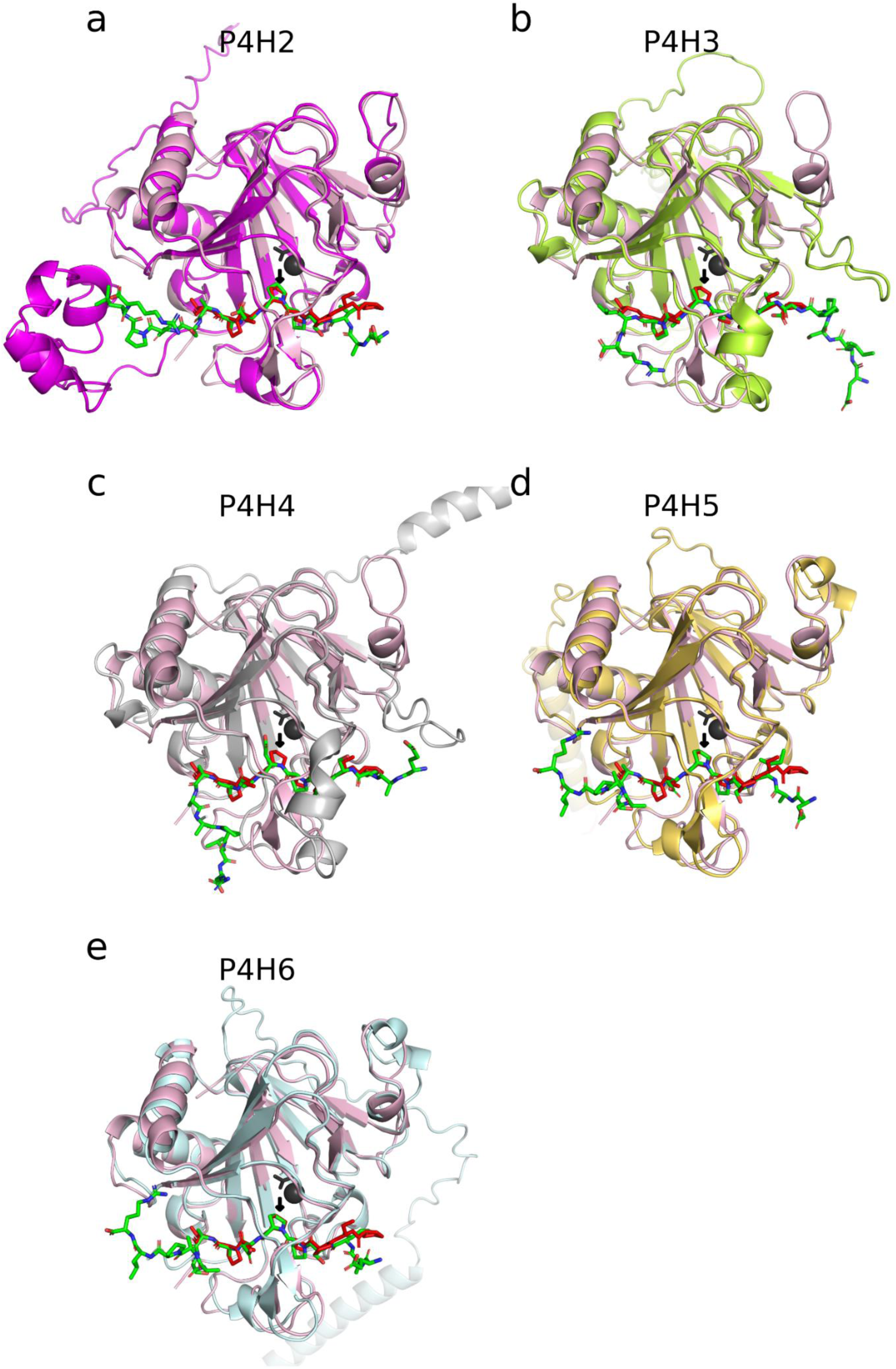
Representative models of the interaction between P4H2-P4H6 and the EPO peptide EAISPPDAASAAPLR. The models of P4H2 (a), P4H3 (b), P4H4 (c), P4H5 (d) and P4H6 (e) with the EPO peptide were generated with AlphaFold2-multimer-v3 without template information. The top-ranking model of the respective P4H and the peptide EAISPPDAASAAPLR (green) were superimposed on the experimentally solved crystal structure of CrP4H1 from *Chlamydomonas reinhardtii* (light red) having a bound (Pro-Ser)_4_ peptide (red) in its active site (PDB ID 3GZE chain C; Koski et al., 2009). The proline from the (Pro-Ser)_4_ peptide within the active site of CrP4H1 where the prolyl-hydroxylation reaction takes place is marked with a black arrow. In the superimposed models of P4H2, P4H5 and P4H6 the second proline of the EPO peptide (EAISP<u>P</u>DAASAAPLR) was located at this position. In P4H3 and P4H4 it was not a proline but an alanine (EAISPPDAAS<u>A</u>APLR) and an aspartic acid (EAISPP<u>D</u>AASAAPLR), respectively.

## Notes

### Competing Interest Statement

The authors have declared no competing interest.

